# QUICHE reveals structural definitions of anti-tumor responses in triple negative breast cancer

**DOI:** 10.1101/2025.01.06.631548

**Authors:** Jolene S. Ranek, Noah F. Greenwald, Mako Goldston, Christine Camacho Fullaway, Cameron Sowers, Alex Kong, Silvana Mouron, Miguel Quintela-Fandino, Robert B. West, Michael Angelo

**Affiliations:** Department of Pathology, Stanford University, Stanford CA, USA; Breast Cancer Clinical Research Unit, Spanish National Cancer Research Center, Madrid, Spain

## Abstract

While recent innovations in spatial biology have driven new insights into how tissue organization is altered in disease, interpreting these datasets in a generalized and scalable fashion remains a challenge. Computational workflows for discovering condition-specific differences in tissue organization typically rely on pairwise comparisons or unsupervised clustering. In many cases, these approaches are computationally expensive, lack statistical rigor, and are insensitive to low-prevalence cellular niches that are nevertheless highly discriminative and predictive of patient outcomes. Here, we present QUICHE – an automated, scalable, and statistically robust method that can be used to discover cellular niches differentially enriched in spatial regions, longitudinal samples, or clinical patient groups. In contrast to existing methods, QUICHE combines local niche detection with interpretable statistical modeling using graph neighborhoods to detect differentially enriched cellular niches, even at low prevalence. Using *in silico* models and spatial proteomic imaging of human tissues, we demonstrate that QUICHE can accurately detect condition-specific cellular niches occurring at a frequency of 0.5% in fewer than 20% of patient samples, outperforming the next best method which required a patient prevalence of 60% for detection. To validate our approach and understand how tumor structure influences recurrence risk in triple negative breast cancer (TNBC), we used QUICHE to comprehensively profile the tumor microenvironment in a multi-center, spatial proteomics cohort consisting of primary surgical resections, analyzing over 2 million cells from 314 patients across 5 medical centers. We discovered cellular niches that were consistently enriched in key regions of the tumor microenvironment, including the tumor-immune border and extracellular matrix remodeling regions, as well as niches statistically-associated with patient outcomes, including recurrence status and recurrence-free survival. The majority of differential niches (74.2%) were specific to patients that did not relapse and formed a robust interconnected network enriched in monocytes, macrophages, APCs, and CD8T cells with tumor and stroma cells. In contrast, the interaction network for patients that relapsed was notably sparse and enriched in B cells, CD68 macrophages and neutrophils. We validated these findings using two independent cohorts, observing similar cellular interactions and predictive power. Collectively, these results suggest that salient, generalized profiles of productive anti-tumor immune responses are defined by a network of structural engagement between innate and adaptive immunity with tumor and stromal cells, rather than by any single specific cell population. We have made QUICHE freely available as a user-friendly open-source Python package at https://github.com/jranek/quiche.

## Introduction

Recent advances in spatial omics technologies [1, 2, 3, 4, 5, 6, 7, 8, 9] have transformed our ability to study how tissue structure, organization, and function changes throughout the course of disease. By simultaneously measuring the molecular profiles and spatial location of individual cells within their native tissue context, these approaches have been used to gain fundamental insights into how cellular interactions influence tissue homeostasis, infectious disease, neurodegeneration, or cancer progression [10, 11, 12, 13, 14, 15, 16, 17, 18, 19]. In cancer, coordinated interactions between heterogeneous cell types within the tumor microenvironment (TME) play a critical role in regulating tumor growth, invasion, metastatic dissemination, and overall treatment effectiveness [20, 21, 22]. For example, the presence of cytotoxic CD8T cells in close spatial proximity to cancer cells has been associated with improved survival and treatment response in patients with triple-negative breast cancer [23, 24], melanoma [25], and colorectal cancer [26]. While advances in spatial profiling technologies have improved our ability to investigate the regulatory mechanisms that drive patient outcomes, analyzing and interpreting these data across heterogeneous patient samples in a generalized and scalable way still presents a significant computational challenge [27].

Computational spatial enrichment methods have been developed to identify cellular niches or tissue architectures from multiplexed images of tissues. These methods can be divided into two general categories: pairwise enrichment methods or spatial clustering approaches. Pairwise enrichment methods [28, 29] test whether two cell types are co-enriched by comparing their observed spatial proximity to what would be expected by random chance using a permutation test. In contrast, spatial clustering approaches [30, 31, 32, 33, 34, 35, 36], otherwise known as microenvironment detection methods, aim to identify higher-order spatial domains within tissue samples. These approaches typically perform unsupervised clustering across all patient samples using a combination of cell spatial coordinates, molecular expression, and/or tissue histology. Computational frameworks, including hidden Markov random fields [37, 35, 36] and graph neural networks [32, 33, 34] are often used to model the spatial relationships between cells and integrate modalities into a consensus representation for clustering. While these approaches can successfully identify cell-cell interactions or tissue structures, they have several limitations when it comes to identifying differential changes in cellular organization across conditions. Namely, they rely on niche prevalence across samples for detection and condition-specific association, which may fail to detect low-prevalence spatial niches that are highly discriminative and predictive of patient outcomes. Moreover, a majority of methods lack robust statistical frameworks for condition-specific testing; as a result, researchers often rely on post-hoc nonparametric statistical tests to assess enrichment [5, 38, 39, 40, 41, 42], which fails to account for the variability in profiled samples limiting niche efficacy.

To address these challenges, we developed QUICHE (QUantitative InterCellular nicHe Enrichment), a statistical framework designed to discover cellular niches differentially-enriched in patient groups, histological structures, or acellular regions. In contrast to previous work, QUICHE combines local niche detection with interpretable statistical modeling using graph neighborhoods to detect differentially enriched cellular niches, even when their prevalence is low. Using *in silico* models and spatial proteomic imaging of human tissues, we show that QUICHE can accurately detect condition-specific cellular niches occurring at low frequency and prevalence, outperforming the next best algorithm by 3-fold in niche recovery by prevalence. To validate our approach and understand how tumor structure influences recurrence risk in triple negative breast cancer (TNBC), we used multiplexed proteomic imaging by time of flight (MIBI-TOF) to comprehensively profile the tumor microenvironment in a multi-center TNBC cohort consisting of 314 primary surgical resections. We applied QUICHE at different spatial resolution scales and identified cellular interactions mediating recurrence-free survival. We validated these findings across two independent cohorts and observed similar cellular interactions and predictive power. Collectively, QUICHE provides a robust and scalable statistical framework for delineating functional cellular interactions within tissue samples that mediate differential outcomes.

## Results

### Overview of the QUICHE algorithm

Here, we present QUICHE – an automated and scalable statistical framework designed to identify differentially-enriched cellular niches in clinical patient groups, histological structures, or acellular regions. Given a multi-sample multiplexed imaging dataset (e.g. MIBI-TOF [1], CODEX [5], IMC [4], Xenium [2]), with the spatial coordinates of cells and their corresponding cell type annotations, QUICHE performs differential spatial enrichment analysis using a three-step process (Figure 1).

**Figure 1:**
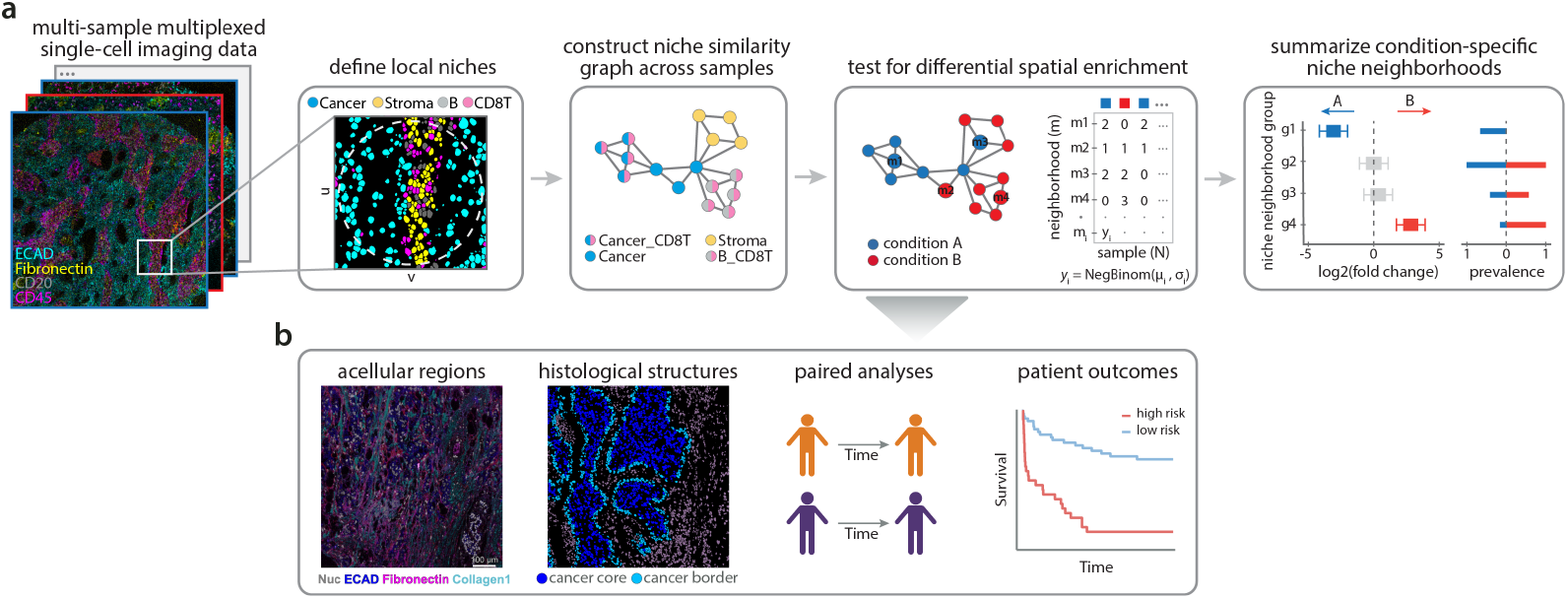
Schematic overview of the QUICHE pipeline. (a) QUICHE performs differential spatial enrichment using a three-step process. In step 1, QUICHE identifies local spatial niches in each sample as the frequency of cell types within a *k*-hop spatial neighborhood bounded by a fixed pixel radius. Niches are then sampled for condition-specific testing using a distribution-focused sketching algorithm that preserves niche frequencies and spectral properties of each sample. In step 2, QUICHE models the similarity of niches across samples using a *k*-nearest neighbor graph, and neighborhood niche counts are modeled using a negative binomial generalized linear model to determine differentially abundant niche neighborhoods. Lastly, in step 3, niche neighborhoods are annotated by the most abundant cell types using a pseudo-bulk approach. (b) QUICHE can be used to discover differentially-enriched cellular niches across spatial regions, longitudinal samples, or clinical patient groups.

In step 1, QUICHE identifies local cellular niches in each tissue sample by analyzing the spatial distribution of cell types. For each cell in a tissue sample, its local niche is defined as the proportion of cell types within *k*-hop spatial neighborhood bounded by a fixed pixel radius. Once local niches are identified, QUICHE performs distribution-focused downsampling [43] to select a subset of representative niches from each sample for condition-specific comparisons. This approach has three main advantages: (1) it controls for any variability in the total number of cells profiled across patients samples, while preserving the underlying distribution of niches, (2) it reduces the number of comparisons, which lowers the multiple hypothesis testing burden and increases statistical power for condition-specific association and (3) it enables scalable analysis of large-scale cohorts.

In step 2, QUICHE tests for differences in the abundance of local cellular niches across conditions using graph neighborhoods, as this approach has shown great success in identifying differentially-enriched cell types [44], cellular expression patterns [45], and quantitative trait loci [46] without relying on unsupervised clustering or pairwise comparisons. QUICHE models the similarity of niches across samples using a *k*-nearest neighbor graph, where the nodes represent niches and the edges describe niche similarity according to pairwise Euclidean distances of cell type frequency vectors. For each neighborhood of the graph, QUICHE counts the number of niches from each patient sample and then tests for differences in the abundance of niches across conditions using a negative binomial generalized linear model (GLM). By leveraging interpretable GLMs, QUICHE can adjust for technical covariates (e.g. batch, study center) or accommodate complex clinical designs when testing for differential enrichment. Finally, statistical significance is determined using a F-statistic, followed by a weighted FDR multiple hypothesis testing correction procedure that accounts for overlapping neighborhoods [47, 44].

Lastly, in step 3, QUICHE annotates niche neighborhoods by the most abundant cell types within them using a pseudobulk based approach for simplified and interpretable downstream analysis. The output of our approach is a set of differentially-enriched niche neighborhoods, their log fold change in enrichment across conditions (e.g. responder/nonresponder, survival time, spatial regions), their statistical significance, and their annotation (Figure 1). By combining local niche detection with interpretable statistical modeling, QUICHE can accurately identify differences in spatial organization across conditions without relying on unsupervised clustering or pairwise comparisons. For a more detailed description of the problem formulation and mathematical foundations behind differential enrichment testing, see QUICHE in the Methods section.

### QUICHE outperforms existing spatial enrichment methods in unstructured and structured spatial topologies

Although pairwise enrichment methods [28, 29] and spatial clustering approaches [48, 39, 49, 41, 50, 30, 31] have successfully identified tissue organizational patterns, there has been no systematic evaluation of whether these approaches can detect differential changes in tissue architecture across patient samples or different experimental conditions. To address this gap and demonstrate the efficacy of QUICHE, we benchmarked five state-of-the-art spatial analysis frameworks (i.e. pairwise enrichment [28], GraphCompass [29], *K*-Means++ [48, 39], CellCharter [30], QUICHE) on their ability to identify condition-specific differences in the spatial organization of cell types using simulated data where the ground truth cellular organization was known (Figure 2a). Each benchmarking study was designed to quantitatively assess how tissue structure, niche size, and niche prevalence across samples influences a method’s ability to detect differential changes in cellular organization. For more details on the simulation design or the spatial enrichment methods evaluated in this study, see *In silico* simulation design or Spatial enrichment approaches in the Methods section, respectively.

**Figure 2:**
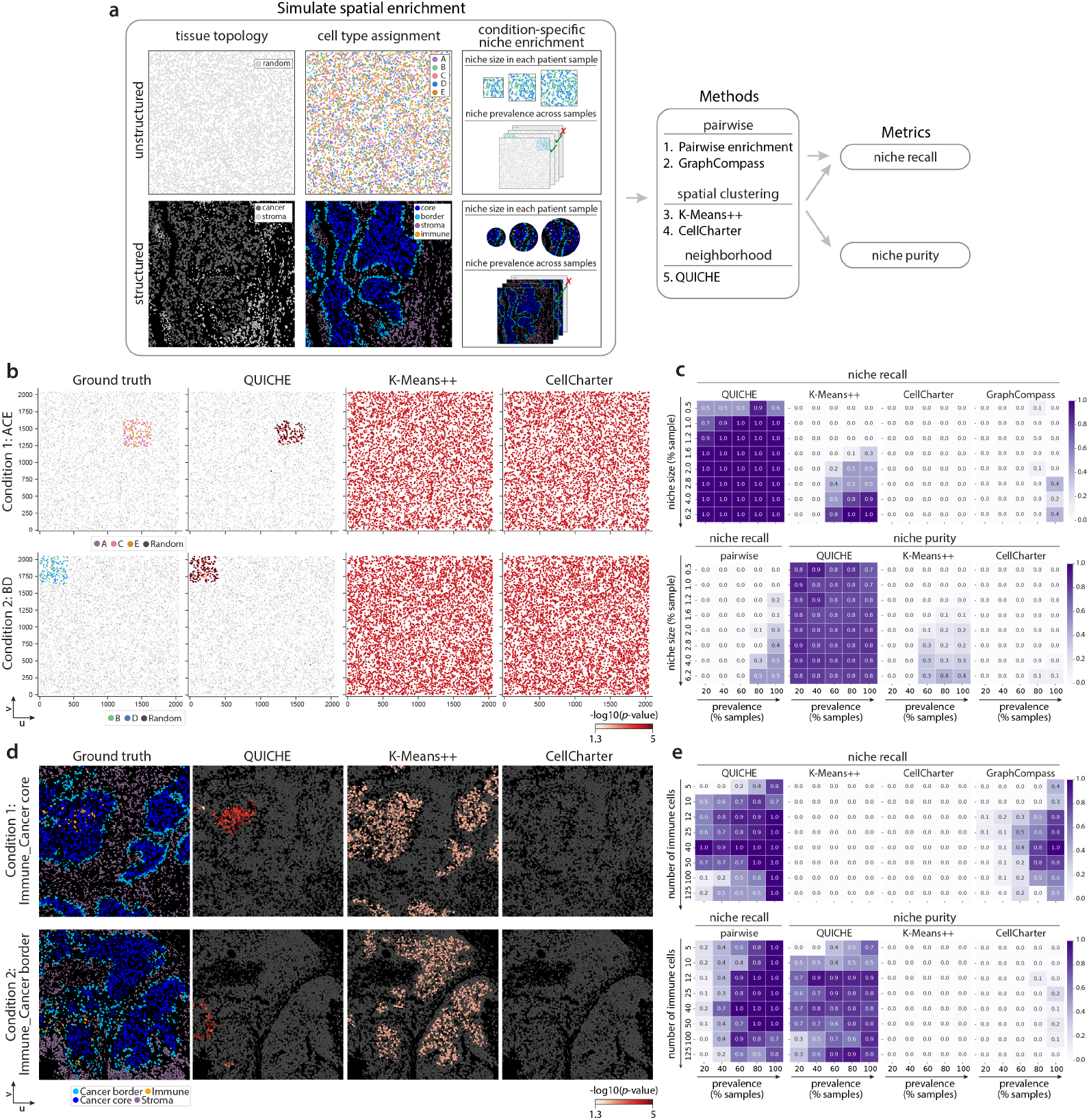
QUICHE discovers differentially-enriched cellular niches across a range of niche sizes, sample prevalence, and tissue structures. (a) Schematic overview of the spatial simulation design. We benchmarked five spatial methods on their ability to recover condition-specific cellular niches while varying the underlying spatial topology, niche size, or sample prevalence. (b) Example visualizations of significant condition-specific cellular niches detected from unstructured spatial topologies (niche size = 4%, niche prevalence = 100%) across different spatial clustering approaches. (c) Performance of five spatial enrichment methods on detecting differentially-enriched cellular niches as measured by niche recall and niche purity. Heatmaps show the average performance across ten simulated unstructured datasets. (d) Example visualizations of significant condition-specific cellular niches detected from structured spatial topologies (niche size = 50 immune cells within 500 pixel radius, niche prevalence = 100%) across different spatial clustering approaches. (e) Performance of five spatial enrichment methods on detecting differentially-enriched cellular niches as measured by niche recall and niche purity. Heatmaps show the average performance across ten simulated structured datasets. *K* = 5 clusters were used for *K*-Means++. Automated cluster detection was used for CellCharter.

In the first simulation study, we generated *in silico* datasets with randomized cell spatial coordinates to evaluate each method’s ability to detect condition-specific cell type interactions in the absence of background tissue organization. Each simulated multi-patient cohort consisted of 20 patient samples, evenly split across two conditions. For each patient sample, we ensured that the number of cells per cell type was identical, and then varied the spatial organization of cell types according to the sample’s condition label. In doing so, this approach allowed us to systematically evaluate each method’s performance solely based on spatial organization. In this simulation design, condition 1 consisted of a niche with cell types A, C, and E spatially co-enriched, whereas condition 2 consisted of a niche with cell types B and D spatially co-enriched. We then investigated the sensitivity of each method to detect condition-specific cellular niches by varying the size of the niche within a sample and its prevalence across samples (See Unstructured simulation in the Methods). For methods that lacked built-in statistical testing for differential enrichment analysis (i.e. *K*-Means++, CellCharter), we performed non-parametric statistical tests between conditions, as this approach is commonly used in the literature [39, 40, 41, 42, 38, 5].

Across all simulated datasets with unstructured spatial topologies, QUICHE successfully identified ground truth cellular niches with high purity and recall (Figure 2b-c). Notably, niches were detected even if they occurred at 0.5% frequency in 20% of patient samples. In contrast, spatial clustering method, *K*-means++ achieved moderate recall scores when niches were large and prevalent; however, often over-partitioned the data resulting in false positives and low purity scores (Figure 2c, Supplementary Figures 1-2). Of note, *K*-Means++ often identified the niche of interest; however, when paired with non-parametric statistical testing, falsely identified alternative condition-specific clusters (e.g. Supplementary Figure 1a). These findings were consistent across different datasets and clustering resolutions (Supplementary Figures 1-2, Supplementary Figure 5a). Lastly, pairwise approaches (i.e. pairwise enrichment, GraphCompass) achieved the smallest sensitivity (Figure 2c), ultimately failing to detect condition-specific differences in unstructured data due to their limited resolution and reliance on a null distribution of shuffled cell types for determining significance.

We then investigated how the underlying tissue structure influences niche detection and condition-specific associations. In this second simulation study, we leveraged spatial proteomic imaging data of compartmentalized tumors to define tumor tissue domains (i.e. cancer core, cancer border, stroma) in each sample (see Structured simulation in the Methods). We then simulated differential immune infiltration within the cancer core (condition 1) or the cancer border (condition 2), and assessed method performance across different niche sizes and sample prevalences. Similar to the unstructured results, QUICHE recovered the majority of ground truth differentially-enriched cellular niches and achieved higher purity scores than the alternative clustering methods (Figure 2d-e). While pairwise approaches (i.e. pairwise enrichment, GraphCompass) successfully identified ground truth condition-specific cellular niches when they were present in a majority of patient samples (60-100% prevalence), they often failed to detect low-prevalence niches. In contrast, spatial clustering approaches (e.g. KMeans++, CellCharter) successfully identified broad tissue domains as intended; however, they often failed to detect finer, condition-specific niches within these domains underscoring the challenges associated with using unsupervised clustering across all patient samples for niche detection and condition-specific association (Supplementary Figures 3-4). These results were consistent across several unsupervised clustering resolution scales (Supplementary Figures 3-5). Importantly, QUICHE was robust to the choice in hyperparameters, and produced reproducible results across a range of hyperparameter choices (See Guidelines on QUICHE parameter selection, Supplementary Table 1, Supplementary Figure 6). Collectively, these results highlight QUICHE’s ability to detect differentially-enriched cellular niches across a range of niche sizes, sample prevalence, and tissue structures.

**Figure 3:**
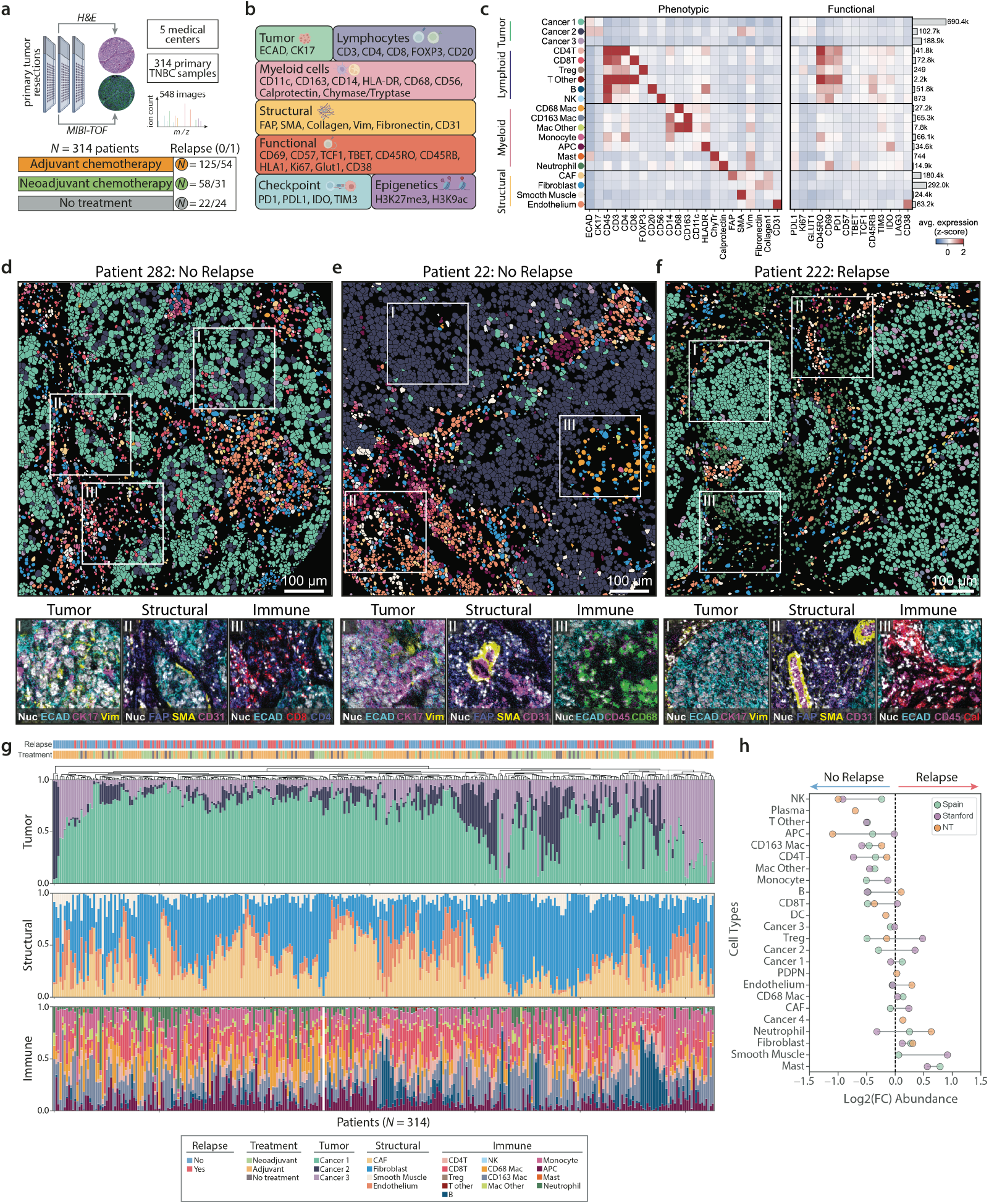
Spatially-resolved single-cell atlas of the primary tumor microenvironment in triple-negative breast cancer. (a) Schematic overview of primary tumor tissue samples profiled with MIBI-TOF imaging. (b) Antibody panel targeting 37 proteins expressed within the primary tumor microenvironment. (c) Heatmap shows the average standardized expression of twenty cell phenotypes clustered according to protein expression of phenotypic markers. Barplots to the right show the total number of cells within each cell phenotype. (d-f) Representative images of TNBC tumors from three patients. Main figure displays the cell phenotype map, while insets show the expression representative protein markers within tumor, structural, and immune regions. (g) Barplots show the frequency of tumor (top), structural (middle) and immune lineages (bottom) across all primary tumor samples in this study. Samples were hierarchically clustered according all frequency subsets. Relapse status and treatment regime are denoted by color in the top row. (h) Scatter plots show the average log2 fold change (FC) in the abundance of cell types across samples from patients that relapsed or did not relapse across three TNBC cohorts.

### Profiling the tumor microenvironment in triple-negative breast cancer

Reasoning that a better understanding of cellular interactions within the primary tumor microenvironment may uncover early risk factors for recurrence, we used multiplexed proteomic imaging by time of flight (MIBI-TOF) [1] to comprehensively profile the tumor microenvironment in a multi-center TNBC cohort consisting of 314 treatment-naive primary surgical resections from the Spanish National Cancer Research Center (CNIO) biobank (Figure 3a, Supplementary Tables 2-3). Among these patients, *N* = 109 experienced a distant recurrence event with a median time to relapse of 479.5 days (Supplementary Table 3). Tumor samples were screened by a breast cancer pathologist to identify regions of interest (up to 4 regions per patient) and tissue microarrays were constructed with cores from each sample for MIBI-TOF imaging (See Methods), using a previously validated 37-plex antibody panel [51, 52] designed to capture various cell types within the TNBC tumor microenvironment, their functional states, and core components of the extracellular matrix and tissue architecture (Figure 3b, TNBC study design and sample collection). This approach resulted in 548 high-plex tissue images of 314 treatment-naive primary TNBC samples (Supplementary Figures 7-9).

To precisely characterize cellular phenotypes within the TNBC tumor microenvironment, we first segmented single cells from multiplexed tissue images using a deep learning algorithm [53] that takes into account nuclear and membrane markers to accurately delineate cells with different morphologies (∼ 2 million total cells with a median of 3479 cells in each of the 548 distinct cores) (Supplementary Figure 10). Following cell segmentation, we performed unsupervised pixel clustering [54] according to phenotypic markers to identify twenty salient cell phenotypes in the tumor microenvironment (Figure 3c). Overall, cancer cells were the most abundant cell population (51.0% total) and consisted of three subsets: Cancer 1 (70.3% of cancer cells), Cancer 2 (10.5% of cancer cells), and Cancer 3 (19.2% of cancer cells) defined by variable expression of epithelial markers (ECAD, CK17), as well as markers for proliferation (Ki67), glucose uptake (GLUT1), and cancer cell invasion and metastatic potential (*α*SMA) [55]. Immune cells were the next most abundant cell type (20.0% total). Activation marker PD-1 was expressed by CD4T (10.9% immune), CD8T (18.9% immune), T other (0.6% immune), Treg (0.1% immune), and B cells (13.4% immune). IDO expression was highest on APCs (9.0% immune) which may suggest long-term immunological tolerance [56]. Expression of CD69 and TIM3 on NK cells (0.2% immune) suggested their maturation, activity, and immunosuppressive status [57, 58]. Lastly, structural cells (29.0% total) included fibroblasts (52.1% structural), cancer-associated fibroblasts [59] (32.2% of structural), endothelial cells (11.3% of structural), and smooth muscle (4.4% of structural). Collectively, cellular phenotypes and abundances were broadly consistent with previous spatial analyses of TNBC (Figure 3c) [51, 16].

We first investigated whether the abundance of particular cellular phenotypes within the primary tumor microenvironment was prognostically predictive of recurrence. Treatment-naive primary TNBC tumors exhibited high inter-patient heterogeneity in the composition of tumor, structural, and immune cell types across recurrence status and treatment regimes (Figure 3d-g). While the abundance of particular cell types was not statistically different between patients who did or did not relapse (Supplementary Figure 13a-b: FDR corrected *p*-values > 0.05), and could not stratify patients by their recurrence status (Supplementary Figure 13c: 5-fold cross validation AUC = 0.522 *±* 0.008), we observed a mild increase in immune infiltration (0 < | log 2(FC) | < 1) in patients who did not relapse (Figure 3h, Supplementary Figure 13, Differential cell type abundance analysis).

To validate these findings and further assess the prognostic impact of cellular composition in the primary tumor microenvironment, we analyzed two independent TNBC cohorts (Supplementary Figures 11-12). For a direct comparison, we used MIBI-TOF imaging to profile the tumor microenvironment in 142 TNBC patients from the Stanford hospital (Supplementary Table 4). Treatment-naive primary tumor resections were profiled using the same antibody panel and phenotyped using the same cell classification schema as our discovery cohort (TNBC study design and sample collection). As a secondary validation, we also analyzed 119 treatment-naive primary tumor samples profiled with imaging mass cytometry from the NeoTRIP chemotherapy trial [60, 61] (NeoTRIP TNBC cohort). Collectively, this approach allowed us to assess the robustness of our findings across different proteomic imaging platforms, antibody panels, clinical cohorts, and medical centers. Across all three cohorts, cell type abundances were not statistically different between relapsers and non-relapsers (Supplementary Figures 13); however, primary tumor samples from non-relapsing patients had consistently higher abundances of CD8T cells, CD4T cells, NK cells, APCs, monocytes, and CD163 macrophages, whereas tumors from patients who relapsed had consistently higher abundances of mast cells and fibroblasts (Figure 3h, Supplementary Figure 13b, e, h). These findings were broadly consistent with previous analyses of TNBC, which also reported limited success in outcome prediction from cell composition in primary tumor samples [60].

To assess the coordination of immune infiltration within the TNBC primary tumor microenvironment, we analyzed the co-occurrence of immune populations across patient samples from all three cohorts (*N* = 575) using a Chi-square test (Immune infiltration analysis). As shown previously [51], we observed a hierarchical ordering of immune populations across patient samples that exhibited immune infiltration (Supplementary Figure 14). For example, all patients that had B cells, also had CD4T cells (*p*-value < 0.05), CD8T cells (*p*-value < 0.05), and macrophages (*p*-value < 0.05). We observed no difference in the interdependence of immune cells in primary tumors from patients who did or did not relapse (Supplementary Figure 14a). These results were consistent across different thresholding parameters and further support a coordinated immune response (Supplementary Figure 14b).

Collectively, these data indicate that recurrence risk in TNBC cannot be determined solely based on the proportion of discrete cell types within primary tumor samples. This highlights the importance of studying the tumor microenvironment beyond cellular densities and underscores the need for spatial profiling technologies and advanced spatial enrichment methods to elucidate the mechanisms of metastatic progression and treatment response.

### QUICHE reveals cellular interactions in the primary tumor microenvironment predictive of recurrence risk

Tumors are spatially-organized ecosystems composed of interacting cell types embedded in an extracellular matrix (ECM), where cells adopt a variety of different functional roles defined by their co-expression of multiple proteins [62]. To understand how cellular interactions within the primary tumor microenvironment influence recurrence risk in TNBC, we performed differential spatial enrichment analysis using QUICHE across three spatial scales: histological structures (e.g. tumor-immune border), acellular regions (e.g. ECM), and local cellular niches across the entire tumor landscape.

Recent studies have suggested that the tumor-immune border may play a prognostic role in TNBC progression [51, 63]. To identify differential immune interactions at this interface, we developed a computational approach to automatically delineate the tumor-immune border (Figure 4a-b). We then used QUICHE to compare the spatial relationships between immune cells at the tumor border as compared to those within the surrounding stroma (See Differentially-enriched celluar niches at the tumor-immune border in the Methods section) (Figure 4c-e). Consistent with previous findings, we observed an enrichment of CD8T cells [64], B cells, and macrophages with cancer cells at the tumor border (Figure 4d). In contrast, CD4T cells were differentially-enriched within the stroma and depleted from the invasive margin in most patient samples (Figure 4e). When examining the functional expression of cell types within differentially-enriched niches, we found that niches within the tumor border consisted of TIM3+ APCs and activated PD1+ CD69+ CD8T cells (Figure 4f), whereas niches within the stroma consisted of IDO+ APCs, memory CD8T cells marked by the expression of CD45RO and memory B cells marked by the expression of CD45RB and CD45RO (Figure 4g). Notably, QUICHE revealed spatial niches with distinct functional expression patterns that were not detectable using traditional cell type abundance approaches (Supplementary Figure 15).

**Figure 4:**
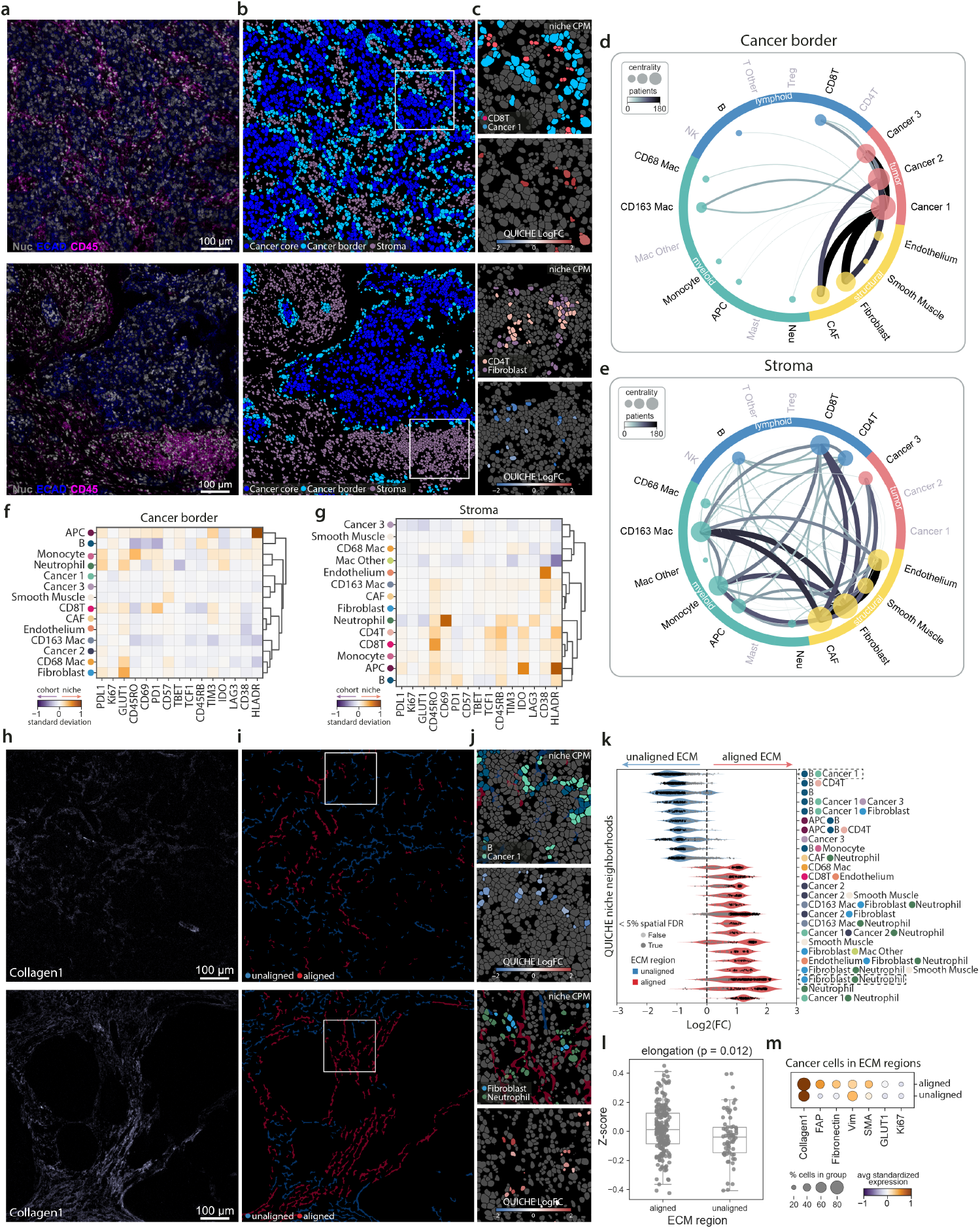
QUICHE identifies differentially-enriched interactions across spatial scales. (a-g) Tumor-immune border interaction analysis. Representative images of tumors annotated by (a) protein expression, (b) tumor regions, or (c) representative cellular niches differentially-enriched within the cancer border (top panel) or stroma (bottom panel). (d) Niche network for cellular niche neighborhoods differentially-enriched at the cancer border. Nodes represent cell types within cancer border-associated niches, and edge weights correspond to the number of unique patients with the corresponding interaction. Node size is proportional to connectedness, as measured by eigenvector centrality. (e) Niche network for cellular niche neighborhoods differentially-enriched in the stroma. (f) Heatmap shows the change in cell type expression within cancer border-associated niche neighborhoods compared to the entire cohort. (g) Heatmap shows the change in cell type expression within stroma-associated niche neighborhoods compared to the entire cohort. (h-m) Extracellular matrix (ECM) alignment analysis. Representative images of tumors annotated by (h) Collagen1 expression, (i) ECM alignment scores, or (j) representative cellular niches differentially-enriched within unaligned (top panel) or aligned (bottom panel) ECM regions. (k) Violin plots show the top 25 differentially-enriched niche neighborhoods within unaligned (blue) or aligned (red) ECM regions. (l) Boxplot shows the average elongation score of cancer cells within aligned or unaligned ECM regions. *p*-values were computed using a two-sided Wilcoxon rank sum test. (m) Dotplot shows the average normalized expression of associated proteins of cancer cells within ECM regions.

We next investigated the role of cellular interactions in remodeling the extracellular matrix and promoting metastatic progression. The extracellular matrix, particularly collagen type I (Collagen1), plays a fundamental role in supporting tumor growth and invasion, where aligned Collagen1 fibers can create structural “highways” that guide cancer cell migration [65, 66]. To examine the relationship between collagen fiber alignment and cellular interactions within the tumor microenvironment, we developed a computational pipeline to automatically detect and segment individual collagen fibers within each tumor sample (Figure 4h-i). After segmenting individual collagen fibers, we devised a fiber alignment score that measures how closely each fiber’s orientation aligns with those of its nearest neighbors (see Differentially-enriched cellular niches in the extracellular matrix in the Methods section). Using this alignment score, we classified collagen regions within each tumor sample as aligned, unaligned, or devoid of ECM (Figure 4i). We then performed differential abundance testing using QUICHE to identify cellular interactions differentially-enriched in aligned versus unaligned collagen regions. Overall, we found that unaligned, randomly oriented collagen regions had greater immune cell infiltration, particularly involving APC, CD4T, and B cell interactions. These regions also supported the formation of tertiary lymphoid structures (niche B–CD4T in Figure 4j-k), which are often associated with sustained local immune responses [67, 68]. In contrast, aligned collagen regions exhibited an immunosuppressive microenvironment enriched in fibroblasts, neutrophils, endothelium, and smooth muscle cells, which may exert mechanical forces on the ECM further promoting the alignment of collagen fibers to create conducive pathways for metastatic spread [65]. Cancer cells within these aligned collagen regions had higher expression of epithelial-to-mesenchymal (EMT) transition markers (e.g. Vimentin (VIM), *α*SMA, fibronectin and FAP) and morphological measurements (e.g. greater elongation, eccentricity, and nucleus-to-cytoplasmic ratio) that support a more migratory phenotype (Figure 4l-m, Supplementary Figure 16). Collectively, these data suggest that unaligned collagen regions may enhance immune surveillance and promote anti-tumor responses, whereas aligned collagen regions may support the recruitment of cells that promote immune evasion and EMT-driven cancer cell migration.

Finally, we aimed to determine whether specific cellular interactions in the primary tumor microenvironment could serve as prognostic indicators of recurrence risk in TNBC. Therefore, we performed statistical differential abundance testing with QUICHE to identify local cellular interactions differentially enriched in primary tumor samples from patients who did or did not experience a recurrence event in the Spain TNBC cohort. More specifically, we used a generalized linear model that included recurrence status and medical center as covariates to ensure that niche enrichment results were not influenced by site-specific factors (See Differentially-enriched cellular niches associated with patient outcomes in the Methods). While cell type abundances were not statistically different between relapsers and non-relapsers (Supplementary Figure 13), QUICHE found notable differences in the spatial organization of cell types that were statistically-associated with recurrence status (mean FDR *p*-value < 0.05) (Figure 5a-k). A majority of differential niches (74.2%) were specific to patients who did not relapse and formed a robust interconnected network consisting of monocytes, macrophages, and T cells, all interacting with the cancer and stroma (Figure 5a, b, d). In contrast, the niche network for patients that relapsed was sparser and enriched for B cells, CD68 macrophages, and neutrophils (Figure 5a, c, e).

**Figure 5:**
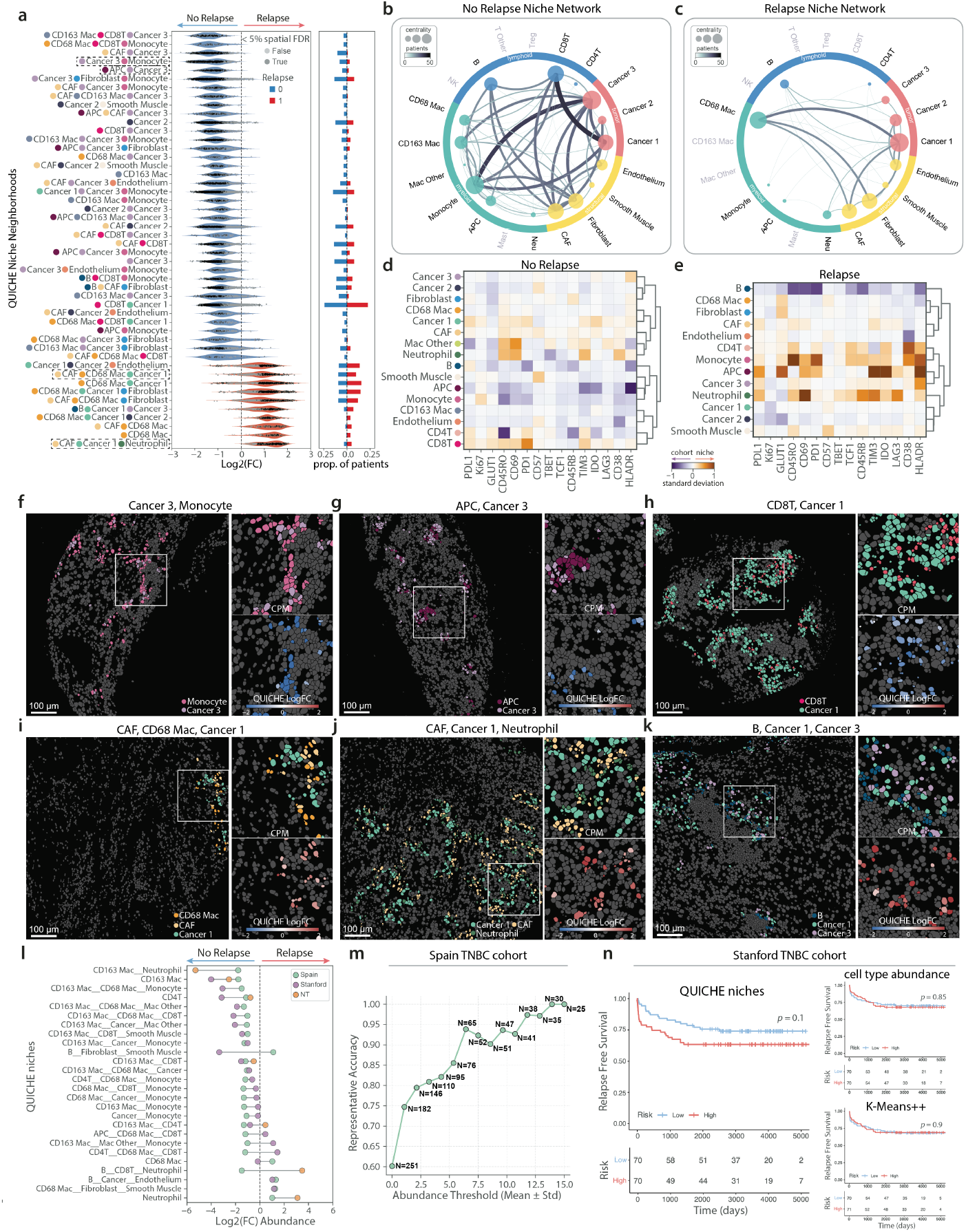
QUICHE reveals cellular niches in the primary TNBC tumor microenvironment predictive of recurrence-risk in TNBC. (a) Violin plots show the top differentially-enriched niche neighborhoods in patients who relapsed (red) or did not relapse (blue). Barplots to the right show the proportion of patients with a niche neighborhood in the respective patient groups. (b) Niche network for non-relapsing patients, where nodes represent cell types within recurrence-free niches and edge weights correspond to the number of unique patients with the corresponding interaction. Node size is proportional to connectedness, as measured by eigenvector centrality. (c) Niche network for relapsing patients, where nodes represent cell types within recurrence-associated niches and edge weights correspond to the number of unique patients with the corresponding interaction. (d) Heatmap shows the change in cell type expression within recurrence-free niche neighborhoods compared to the entire cohort. (e) Heatmap shows the change in cell type expression within recurrence-associated niche neighborhoods compared to the entire cohort. (f-k) Representative images of six differentially-enriched niche neighborhoods. Insets show cells within each niche neighborhood colored according to cell type (top) or predicted enrichment score (bottom). (l) Scatter plots show the average log2 fold change in cell type abundance between patients who did or did not relapse across three cohorts. (m) Line plot shows the representative accuracy of QUICHE niche neighborhoods in the Spain cohort according to the ratio of positive to negative niches over several abundance cutoffs. (n) Kaplan-Meier curves showing relapse-free survival (y-axis) as a function of time (x-axis). A Cox proportional hazards model was trained on the abundance of QUICHE niche neighborhoods (left), cell type abundance (right top), or *K*-Means++ spatial clusters (right bottom) in the Spain TNBC cohort to predict recurrence risk in gthe Stanford TNBC cohort. Patients were stratified into two groups based on the median risk score. *p*-values were calculated using a log-rank test to compare recurrence free survival differences between high- and low-risk groups.

Collectively, these results suggest that productive anti-tumor responses are defined by both innate and adaptive immunity rather than a single specific cell population. In contrast, rather than being defined by what they contain, we found that primary tumor samples from patients who later developed recurrence were most notable for what they lacked. High risk lesions had less recurring sets of cellular interactions across patient samples, which may highlight the challenges associated with identifying universal biomarkers for treatment. We evaluated the representative accuracy of QUICHE niche neighborhoods in classifying recurrence risk from the Spain TNBC cohort by computing the ratio of recurrence-associated to recurrence-free niches within each patient sample over a range of abundance thresholds (See Differentially-enriched cellular niches associated with patient outcomes). Overall, we found that increasing the niche abundance threshold by 2.5 standard deviations from the mean could classify 100 patients with 90% accuracy (Figure 5m).

One of the main advantages of QUICHE is in its ability to assess how cells are functionally interacting within outcome-associated niches as compared to the entire cohort. For example, we found that non-relapsing patients had cellular niches with naive B cells marked by the lack of expression of CD45RO and CD45RB, activated PD1+ CD8T cells, HLADR+ Cancer 3 cells which has been associated with good prognosis [69, 60], and proliferating Ki67+ monocytes (Figure 5d). In contrast, although the relapse-associated niche network was notably sparser and had less recurring sets of cellular interactions across patients, cells within significant relapse-associated niches had fundamentally distinct expression profiles. For example, niches contained CD69+ neutrophils, PD1+ monocytes, TIM3+ IDO+ APCs, and naive TCF1+ CD45RB+ CD45RO+ CD4T cells (Figure 5e). Notably, these phenotypes were not observable when examining the functional expression profiles of cells across the entire set of relapsers or non-relapsers (Supplementary Figure 17). Taken together, these results highlight how the structural organization of cells within primary tumor samples dictate their function and overall phenotype and highlight the importance using a spatial enrichment method for detecting local cellular niches across different subsets of patients.

We validated these findings across two independent TNBC cohorts and found similar sets of cellular interactions (Figure 5i, Supplementary Figures 11e-g, 12e-g). Notably, patients who did not experience a relapse event showed increased enrichment of CD163 macrophage interactions, CD8T–cancer interactions, and monocyte–cancer interactions. In contrast, patients that relapsed had differential enrichment of B– cancer interactions and neutrophils (Figure 5i). To test whether cellular interactions within the primary tumor microenvironment were independently predictive of recurrence, we trained a Cox proportional hazards model [70] on the abundance of QUICHE niches in each patient sample from our discovery TNBC cohort (Cox proportional hazards model). We then applied this trained model to predict recurrence risk in our independent TNBC cohort (142 patients, 277 images, ∼1 million total cells) profiled with the same antibody panel and phenotyped with the same cell classification schema. Notably, QUICHE niches independently stratified patients by recurrence risk (log-rank test *p*-value = 0.1), which was not possible using traditional cell type abundances (log-rank test *p*-value = 0.85) or unsupervised microenvironment detection with *K*-Means++ clustering (log-rank test *p*-value = 0.9) (Figure 5n).

Collectively, these results highlight the prognostic power of recurrence-associated QUICHE niches and demonstrate the model’s robustness in uncovering functionally relevant cellular interactions and spatial organization patterns linked to patient outcomes across independent cohorts.

## Discussion

Spatial enrichment methods have transformed our ability to investigate how tissue organization changes in health and disease; however, existing methods are limited by (1) an inability to detect localized cellular niches across spatial scales and (2) a lack of robust statistical frameworks for condition-specific testing. To overcome these limitations, we developed QUICHE, a scalable framework that leverages graph neighborhoods and interpretable statistical modeling to detect differentially enriched cellular niches without relying on pairwise cell type comparisons or pre-existing sets of clusters. The key innovation of this approach is in its ability to detect low-prevalence spatial niches that are differentially enriched across conditions while accounting for sample variability, thereby enabling a more robust and scalable analysis of complex tissue architectures across large-scale clinical cohorts.

In this study, we benchmarked five state-of-the-art spatial methods on their ability to identify condition-specific differences in cellular organization using *in silico* data where the ground truth cellular organization was known. Through a series of quantitative comparisons, we illustrate how niche size, niche prevalence, and underlying spatial topology can considerably impact the performance of existing spatial methods on identifying condition-specific differences in cellular organization. On unstructured data, we found QUICHE achieved significant improvement in niche recovery and niche purity, as compared to the traditional pairwise enrichment and spatial clustering approaches. This improved performance can be attributed to QUICHE’s niche resolution and its ability to detect niches without relying on comparisons to randomly shuffled distributions of cells or nonparametric statistical tests. In contrast, on datasets with structured spatial topologies, QUICHE outperformed methods in its ability to accurately resolve complex niche interactions (e.g. immune cells) within well-defined tissue domains (e.g. tumor core, tumor border). In this context, unsupervised clustering approaches could successfully identify background tumor structure; however, either failed to detect small condition-specific differences in cellular organization or over-partitioned the data resulting in numerous false positives depending on the clustering resolution. These findings highlight the importance of considering underlying spatial structure, niche size, and sample prevalence when performing differential enrichment analyses.

To investigate how tumor structure influences recurrence risk in TNBC, we used MIBI-TOF imaging to profile treatment-naive primary tumor samples from a total of 456 TNBC patients spanning two clinical cohorts and six medical centers. We then used QUICHE to perform an in-depth spatially-resolved single-cell analysis of the TNBC tumor microenvironment, analyzing cellular interactions that could serve as prognostic indicators of recurrence risk, as well as those that may be mediating anti-tumor responses or metastastic progression within the tumor-immune border or extracellular matrix. While cell type abundance was insufficient to prognostically predict recurrence risk, our analyses revealed significant differences in the cellular organization and functional role of cell types that stratified patient outcomes. Specifically, non-relapsing patients exhibited a redundant network of innate and adaptive immune responses involving monocytes, macrophages, and T cell subsets interacting with tumor and stromal cells. In contrast, relapsing patients, despite having a similar degree of immune infiltration, had less re-occurring sets of interactions across patient samples and was enriched for CD68 macrophage and B cell interactions. Spatial cellular niches were validated across two independent TNBC cohorts and could stratify recurrence risk; this result was not attainable using cell type abundance or the alternative spatial clustering methods. Collectively these findings underscore the need for robust spatial enrichment methods to effectively analyze heterogeneous responses in large-scale clinical cohorts, and demonstrate the overall robustness and generalizability of QUICHE.

Overall, this study highlights how spatial profiling technologies, local niche detection, and interpretable statistical modeling can be used to uncover cells that may be functionally interacting in tissues leading to differential outcomes. However, several limitations should be considered prior to performing spatial enrichment analysis. First, while QUICHE can identify differentially-enriched cellular niches, its performance relies heavily on the quality of cell segmentation and cell annotations used as input; thus, quality control preprocessing or batch effect correction should be performed prior to spatial enrichment analysis. Second, in this study, we provided a framework for spatial analysis using a fixed spatial resolution (e.g. 200 pixels), which may not capture hierarchical or multi-scale interactions in tissues. Third, QUICHE relies on a *k*-nearest neighbor graph for condition-specific testing, which may fail to identify rare niches that are represented by a small number of neighborhoods. In this scenario, we recommend choosing a smaller *k* or increasing the subsampling size. Lastly, for applications focused on analyzing the spatial distribution of rare cell types, it may not be appropriate to label niche neighborhoods according to their most abundant cell types. Therefore, we recommend expanding niche neighborhood annotations to more cell types, or examining those niches individually. Future work could focus on extending this framework (1) across spatial resolution scales to uncover hierarchical multi-scale cellular interactions or (2) across multi-modal spatial data to uncover the interplay between cellular phenotypes and functional states. As the amount of multiplexed imaging data continues to grow, it is crucial we develop automated and scalable tools that can effectively translate heterogeneous datasets into actionable insights. In this context, QUICHE offers a flexible and scalable solution for differential spatial analysis.

## Methods

### QUICHE

#### Problem formulation

Let 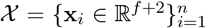 denote a multiplexed imaging dataset (e.g. MIBI), where **x**_*i*_ represents the vector of *f* features (e.g. proteins) measured in cell *i* with (*u*_*i*_, *v*_*i*_) spatial coordinates in the tissue sample. Given a multi-patient cohort, 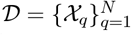 consisting of *N* patient samples, where each patient sample, *q*, has a clinical attribute label, *g*_*q*_ ∈ {1, …, *G*} (e.g. relapse, no relapse), our goal is to build a statistical model to identify differences in the spatial organization of cell types across experimental conditions to predict patient-level clinical outcomes, *g*. Such an approach is poised to uncover how cell types functionally interact in individual patient samples, illuminating the mechanisms behind why patients with similar clinical characteristics respond differently to treatment.

#### Defining local cellular niches

To classify the spatial organization of cells amongst patients with different conditions or clinical outcomes, we first define local cellular niches in each patient sample as the proportion of cell types within a *k*-hop spatial neighborhood bounded by a fixed pixel radius, *s*, around each cell (e.g. *s* = 200 pixels). That is, for cell *i* in sample *q* with *T* cell types, we define its local niche as the proportion, 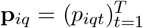, where 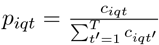 and *t* ∈ {1, …, *T*}. Here, *c*_*iqt*_ represents the number of cells of type *t* within cell *i*’s spatial cellular neighborhood, 𝒩_*iq*_ ={*j*: *j* ∈ 𝒱, (*i, j*) ∈ *ε*, and|| (*u*_*i*_, *v*_*i*_) − (*u*_*j*_, *v*_*j*_) || ≤ *s*}. Each niche **p**_*iq*_ corresponds to a single index cell in a given patient sample and captures the local distribution and spatial organization of cell types. Niches are defined for every cell in a sample. We then perform distribution-focused downsampling [43] to select a subset of representative niches, *l << n*, from each patient sample for downstream condition-specific comparisons. In doing so, this downsampling approach 1) controls for any variability in the total number of profiled cells per patient sample, while preserving the underlying distribution of niches; 2) reduces the number of niches that need to be compared in multiple hypothesis testing, increasing the statistical power; and 3) aids in the scalability of analysis.

#### Differential spatial enrichment

To test for differential spatial organization of cell types across conditions, we propose to model differences in the abundance of local cellular niches using graph neighborhoods, as this approach has shown great success in identifying cell types [44], cellular expression patterns [45], and quantitative trait loci [46] statistically-associated with experimental conditions without relying on unsupervised clustering or pairwise comparisons. Specifically, we model the similarity of niches across patient samples using a *k*-nearest neighbor graph, 𝒢 = (𝒱, ε), where the nodes 𝒱 represent niches and the edges are defined according to pairwise Euclidean distances of cell type frequency vectors, **p**_*iq*_. For each niche neighborhood in the graph, *m* where *m* ∈ {1, …, *l*}, we count the number of niches from each patient sample to construct a neighborhood by sample, *m* × *N*, count matrix. We then test for differences in the abundance across conditions using a negative binomial generalized linear model as, 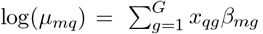, where *y*_*mq*_ represents the observed count of niches from patient *q* in neighborhood *m, µ*_*mq*_ is the mean number of niches from patient *q* in neighborhood *m*, such that *y*_*mq*_ ∼ NegBinom(*µ*_*mq*_, *ϕ*_*m*_), and *x*_*qg*_ is an indicator vector of condition *g*. Here, *β*_*mg*_ is the regression coefficient indicating the log fold change in enrichment of a niche neighborhood in condition *g* (e.g. relapse) as compared to all other conditions (e.g. no relapse). Lastly, we perform hypothesis testing using a F-statistic with a weighted FDR multiple hypothesis testing correction procedure as introduced in Refs. [44, 47]. This approach accounts for overlapping neighborhoods by adjusting *p*-values according to the connectivity of the graph. Therefore, the output of our approach is a set of spatially reoccurring cellular niches that are statistically-associated with patient outcomes, *g*. Notably, by testing for differential spatial enrichment on graph neighborhoods, QUICHE can identify local cellular niches associated with patient outcomes across different scales of patient prevalence.

#### Niche annotations

We annotated niche neighborhoods according to the average cell type abundance within each niche neighborhood as,

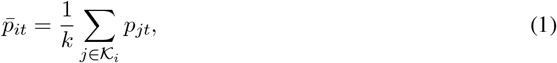

where 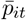 is the average abundance of cell type *t* in the neighborhood of niche *i, p*_*jt*_ is the proportion of cell type *t* in niche *j*, and 𝒦_*i*_ is the set of *k*-most similar niches around niche *i* based on the *k*-nearest neighbor graph. We then labeled each niche neighborhood according to the most abundant cell types in the averaged vector 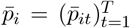. Cell types within niche neighborhood groups were labeled in alphabetical order for simplicity.

#### Niche networks

To visualize the interactions between cell types within significant niches, we constructed condition-specific niche networks, 𝒫^*g*^ = (𝒱^*g*^, ε^*g*^), where the nodes 𝒱^*g*^ represent cell types and the edges *ε*^*g*^ connect cell types that co-occur within significant niches across patients under condition *g*. Specifically, the edge weight *w*_*ij*_ between cell types *i* and *j* quantifies strength of association as measured by the number of unique patients that have the cellular interaction from the set of significant niches as,

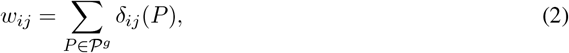

where 𝒫^*g*^ is the set of patients under condition *g* and *δ*_*ij*_(*P*) is an indicator function defined by,

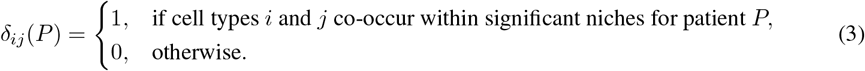

### Guidelines on QUICHE parameter selection

In this section, we provide practical recommendations for selecting hyperparameters in QUICHE (Supplementary Table 1) and describe QUICHE’s sensitivity to choices in hyperparameters for simulated datasets with structured spatial topologies. We recommend standard quality control preprocessing prior to performing spatial enrichment analysis with QUICHE.

#### Number of neighbors in spatial niche detection (n_neighbors)

QUICHE determines local cellular niches in each patient sample as the proportion of cell types within a *k*-hop spatial neighborhood bounded by a fixed pixel radius. The choice of number of hops or the number of neighbors directly influences the definition of the niche frequency vector. We recommend choosing a neighbor size (n_neighbors) to account for the complexity of expected niches.

#### Size of radius in spatial niche detection (radius)

The radius size (measured in pixels) defines the outer boundary when identifying local cellular niches in each patient sample. We recommend selecting a radius size based on the spatial scale of cellular interactions of interest. For interactions with larger spatial topologies (e.g. ducts), we recommend performing enrichment analysis using all cells within a defined radius.

#### Number of sampled niches from each patient sample (sketch size)

To aid in the scalability of analysis of large clinical cohorts and reduce the multiple hypothesis testing burden, QUICHE samples a subset of niches from each patient sample for condition-specific comparisons. We have previously shown that selecting a small representative subset of cells with Kernel Herding sketching [43, 71] preserves the original distribution of cell states, (2) achieves the same fidelity of downstream performance with far fewer cells, and (3) results in improved scalability and faster runtimes. However, it is important to note that selecting too small of a sketch size (i.e. too few subsampled niches, *l* << *n*) has the potential to neglect rare niches, which may impact condition-specific niche enrichment. Therefore, we recommend selecting a larger number of subsampled niches from the original dataset to retain the expected rare niches that occur at a small frequency.

#### Number of nearest neighbors in niche similarity graph construction (k_sim)

QUICHE uses a *k*-nearest neighbor graph to test for differences in the abundance of cellular niche neighborhoods across conditions. The choice in the number of nearest neighbors, *k*_sim, directly influences the connectivity of the graph. Selecting a small value of *k*_sim will connect more similar niches together and preserve niche states; however, it also reduces the number of niche counts per patient sample per condition. Selecting a large value of *k* can increase power to detect differences; however, this will also increase the multiple hypothesis testing burden. Overall, we recommend choosing (*k*_sim = 50 − 300) depending on the number of cells and number of patient samples. We recommend selecting a smaller *k* (e.g. *k*_sim = 50) when trying to identify rare niche neighborhoods.

#### QUICHE’s sensitivity and robustness to hyperparameters

To evaluate QUICHE’s sensitivity to choices in hyperparameters, we varied the number of neighbors in spatial niche identification (n_neighbors = 10, 20, 30, 40, 50, 75, 100), the radius size in spatial niche detection (radius = 50, 75, 100, 200, 250, 300 pixels), the number of sampled niches from each patient sample (sketch size = 250, 500, 750, 1000, 2000, 2500, total), and the number of nearest neighbors in niche similarity graph construction (*k*_sim = 50, 100, 200, 300). For each parameter configuration, we fixed all of the remaining parameters to be the same (n_neighbors = 30, radius = 200, sketch size = total, and *k*_sim = 100), and then varied the parameter of interest. Here, we assessed performance on structured simulated datasets (niche size = 50 immune cells in radius 500 and niche prevalence = 100% patient samples) by computing the recall and purity of predicted niches to the ground truth across 5 random trials for each dataset and parameter configuration. Overall, we found QUICHE was robust to choices in hyperparameters (Supplementary Figure 6). However, we note that parameter choices should be selected according to the dataset of interest.

### Spatial enrichment approaches

#### Pairwise Enrichment

Pairwise cell type enrichment scores [28] were calculated for all possible pairs of cell types in each patient sample. For each cell type pair, the observed co-enrichment was determined by constructing a spatial neighbor graph, where the nodes represent cells and the edges connect similar cells according to spatial proximity using Delaunay triangulation [72]. Enrichment scores were then computed by comparing the observed co-enrichment to a null distribution using a Z-score. To evaluate condition-specific differences, pairwise enrichment scores were compared between groups using a two-sided Wilcoxon rank sum test using the `ranksums` function in Scipy v1.12.0. For our subsequent analyses, we considered a cell type pair to be significant if the *p*-value < 0.05.

#### GraphCompass

GraphCompass [29] is a computational framework designed for assessing differences in spatial organization across conditions. For cellular niche analysis, GraphCompass first performs pairwise cell type enrichment as described above, where cell type proximity is determined using Delaunay triangulation of cell spatial coordinates [72]. Next, GraphCompass fits a linear model to the pairwise enrichment *Z*-scores, with a fixed linear term to account for the patient covariate and an interaction term between all levels of the condition and cell type pairs. The model then performs a *t*-test to determine if a coefficient is significant. For our subsequent analyses, we considered a cell type pair to be significant if the *p*-value < 0.05.

*K***-means++** *K*-Means++ [73] is an unsupervised learning algorithm that partitions data points into *K* clusters by iteratively minimizing the Euclidean distance between data points and their nearest centroid until convergence. To identify spatial clusters, we implemented *K*-means++ clustering as previously described [39, 50]. First, local cellular niches were defined in each patient sample as the proportion of cell types within a fixed pixel radius around each cell (e.g. *s* = 200 pixels). That is, for cell *i* in sample *q* with *T* cell types, we defined its local niche as the proportion, 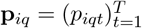, where 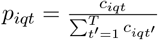 and *t* ∈ {1, …, *T*}. Here, *c*_*iqt*_ represents the number of cells of type *t* within cell *i*’s spatial cellular neighborhood, 𝒩_*iq*_ = {*j*: ||(*u*_*i*_, *v*_*i*_) − (*u*_*j*_, *v*_*j*_)|| ≤ *s*}. Next, we vertically concatenated cell type proportion vectors across patient samples to construct a n * N × cell type matrix. We then performed *K*-Means++ clustering (*K* = 3, 5, 7) to identify spatial clusters using the `KMeans` function in scikit-learn v1.5.2 in Python. To test for condition-specific differences, we computed the proportion of spatial clusters in each patient sample and then tested for differences between patient groups using a Wilcoxon rank sum test with the `ranksums` function in Scipy v1.12.0. For our subsequent analyses, we considered a spatial cluster to be significant if the *p*-value < 0.05.

#### CellCharter

CellCharter [74] is a spatial analysis framework that identifies cellular niches or spatial clusters in a four step process. First, a spatial graph is constructed, where the nodes represent cells and the edges connect similar cells according to spatial proximity using Delaunay triangulation. Next, CellCharter transforms the expression data into a latent subspace representation, *Z*, using dimensionality reduction and batch effect correction. For each cell, feature vectors are then aggregated across neighbors up to a defined layer *l* ∈ {0, …, *L*} using aggregation functions (e.g. mean, standard deviation, min, max). More specifically, let *A*^(*l*)^ represent the adjacency matrix of neighbors at layer *l, f*_*j*_ for *j* ∈ {0, …, *J*} represent the aggregation functions, (*X, Y*) represent the matrix concatenation operation, and *XY* represent the matrix inner product operation. The output of the neighborhood aggregation step is

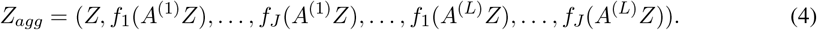

Cells are then clustered into microenvironments using a Gaussian mixture model on *Z*_*agg*_, where the number of microenvironments are chosen according to the most stable assignment using the Fowlkes-Mallows Index [75]. Given that our *in silco* data were not simulated with batch effects, we performed dimensionality reduction using principal components analysis (PCs = 50) [76, 77]. Neighborhood aggregation (*l* = 3, aggregation function = mean) and clustering parameters (runs = 10) were chosen according to default parameters. To test for condition-specific differences, we computed the proportion of microenvironments in each patient sample and then tested for differences between patient groups using a Wilcoxon rank sum test with the `ranksums` function in Scipy v1.12.0. For our subsequent analyses, we considered a microenvironment to be significant if the *p*-value < 0.05. CellCharter was implemented using the cellcharter v0.3.1 package in Python across a range of clustering resolutions (*K* = 3, 5, 7, auto-selected).

#### QUICHE

QUICHE was run as previously described. For our simulation tests, niches were determined using a spatial proximity graph of 30 neighbors bounded by a fixed 200 pixel radius. A niche similarity graph was constructed using a *k*-nearest neighbor graph (*k* = 100), and niche neighborhoods were considered significant if *p*-value < 0.05.

#### *In silico* simulation design

To validate our approach and benchmark spatial enrichment methods on their ability to identify condition-specific differences in the spatial organization of cell types from individual patient samples, we simulated multi-patient cohorts with different spatial topologies (e.g. unstructured, structured), where the ground truth organization was known. Here, each multi-patient cohort consisted of 20 patient samples evenly split across two conditions. For each patient sample, we fixed the number of cells and cell types to be the same, and then varied the spatial organization of cells according to each patient’s condition label (e.g. condition A consisted of immune cells enriched in the cancer core).

#### Unstructured simulation

We generated simulated datasets with unstructured spatial topologies using a two-step approach. First, for each patient sample, we randomly sampled the spatial coordinates of 5000 cells across five cell types from a uniform distribution within a 2048 × 2048 pixel domain. Each image was then discretized into a *g* × *g* grid, and one grid was randomly selected for condition-specific niche enrichment. Here, the grid size, *g*, is directly proportional to the niche prevalence (i.e. 5 × 5 grid indicates that a niche occupies 1*/*25 or 4% of a patient sample). The output of this simulation approach is a ground truth reference consisting of cell-to-cell type and cell-to-niche assignments. To evaluate how well spatial enrichment methods can identify condition-specific differences in the spatial organization of cell types with unstructured spatial topologies, we varied both the size of a niche in a patient sample (i.e. grid size = 3, 4, 5, 6, 7, 8, 9, 10) and its prevalence across patient samples (i.e. 20, 40, 60, 80, 100% of patients have the niche). Method performance was evaluated over five simulated datasets for each parameter configuration.

#### Structured simulation

For a more realistic simulation design, we leveraged multiplexed proteomic imaging data from the Spain TNBC cohort to define cell spatial coordinates and tissue domains as follows.

##### Tumor selection

To generate simulated datasets with structural spatial topologies, we randomly selected tumors with compartmentalized tumor-immune spatial architectures using the mixing score defined in Ref. [51] as,

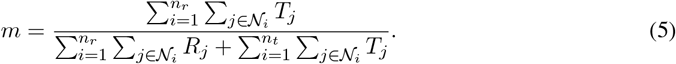

The mixing score quantifies the degree of mixing between reference (i.e. tumor) and target (i.e. immune) cells within a fixed spatial radius, *s*, by computing the ratio of heterotypic (e.g. cancer-immune) to homotypic interactions (e.g. cancer-cancer, immune-immune). Here, *n*_*r*_ and *n*_*t*_ represent the number of reference and target cells, respectively. The indicator functions *T*_*j*_ and *R*_*j*_ denote whether a target or reference cell type is within the neighborhood 𝒩 of cell *i*. Specifically, *T*_*j*_ = 1 if cell *j* is a target cell type within the neighborhood 𝒩_*i*_, and *T*_*j*_ = 0 otherwise. Similarly, *R*_*j*_ = 1 if cell *j* is a reference cell type within the neighborhood 𝒩_*i*_, and *R*_*j*_ = 0 otherwise. The neighborhood 𝒩_*i*_ is defined as {*j*: ‖ (*u*_*i*_, *v*_*i*_) − (*u*_*j*_, *v*_*j*_) ‖ ≤ *s*}, where (*u*_*i*_, *v*_*i*_) and (*u*_*j*_, *v*_*j*_) are the spatial coordinates of cells *i* and *j*, respectively. Tumors were considered to have compartmentalized spatial architectures if they had more homotypic than heterotypic interactions.

##### Tumor compartments

To define structural regions in each tumor image, we followed the approach outlined previously [52]. Briefly, tumor compartment regions (i.e. cancer core, cancer border, stroma) were defined by taking the union of the ECAD signal with the cancer cell segmentation masks. Tumor compartment masks were then thresholded (0.0015), binarized, and eroded by 100 pixels to define cancer cells within the core and border. All remaining cells were defined as stroma. Image processing was performed using the scikit-image v0.19.3 package in Python.

##### Differential spatial enrichment

We induced condition-specific cell type enrichment within structured spatial topologies using a two-step approach. First, for each patient sample, we randomly sampled cells from condition-specific tumor regions (e.g. cancer core, cancer border) using a Gaussian distribution. Next, a subset of cells within a specified radius (radius = 50, 100, 250, 500) were reclassified as immune cells to create localized clusters of immune infiltration within tumor regions. We then evaluated how well spatial enrichment methods can detect condition-specific differences in cellular organization by varying the (1) niche size (radius = 100, 250, 500 pixels), (2) niche sparsity (5, 10, 25% of immune cells within the radius), and (3) niche prevalence across patient samples (20, 40, 60, 80, 100% of patients have the niche). Method performance was evaluated over five simulated datasets for each parameter configuration.

##### Expression

Splatter [78] is a single-cell RNA sequencing simulation software that generates count data using a gamma-Poisson hierarchical model, with modifications to alter the mean-variance relationship amongst genes, the library size, or sparsity. We used Splatter to simulate expression data in our *in silico* multi-patient cohorts using in a two-step process. First, simulation parameters were estimated by fitting the model to a real single-cell RNA sequencing dataset [79]. Next, the estimated parameters (mean_rate = 0.0173, mean_shape = 0.54, lib_loc = 12.6, lib_scale = 0.423, out_prob = 0.000342, out_fac_loc = 0.1, out_fac_scale = 0.4, bcv = 0.1, bcv_df = 90.2, dropout = None) were used in the Splatter groups function (python wrapper scprep SplatSimulate v1.2.3 of splatter v1.18.2) to simulate expression data (*p* = 500) for each cell type (e.g. cancer core, cancer border, stroma, immune).

#### Method Evaluation

To quantify differential spatial enrichment performance, we compared the predicted niches to the ground truth annotations using two metrics, niche recall and niche purity. Given that each method returns a different output (e.g. clusters, pairwise associations, index cell-level scores), we employed a diverse set of metrics to quantify overall performance. Both niche recall and niche purity return a score between 0 and 1, where 1 indicates that the predicted labels were in perfect accordance with the ground truth annotations.

##### Niche Recall

To assess a method’s ability to recover the ground truth niches, we computed a niche recall score as,

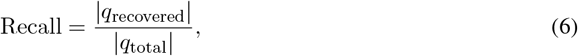

where |*q*_recovered_| indicates the number of ground truth niches recovered and |*q*_total_| indicates the total number of ground truth niches. For the pairwise enrichment methods (i.e. pairwise enrichment, GraphCompass), a niche was considered recovered if all possible pairwise combinations of cell types were significantly enriched (e.g. niche ACE required AC, CE, and AE). For the spatial microenvironment detection methods (e.g. *K*-Means++, CellCharter, QUICHE) a niche was considered recovered if a significant predicted cluster had a Purity_*c*_ ≥ 0.5.

##### Niche Purity

To quantify the degree of intermixing in predicted spatial microenvironment clusters, we computed a niche purity score as,

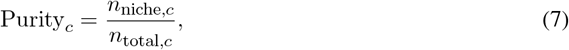

where *n*_niche,*c*_ indicates the number of ground truth niche cells in predicted cluster *c* and *n*_total,*c*_ indicates the total number of cells in predicted cluster *c*.

#### Spain and Stanford TNBC study design and sample collection

To investigate how the tumor microenvironment contributes to recurrence risk in TNBC, we performed a retrospective study analyzing treatment-naive primary tumor samples from two independent cohorts, the Spain TNBC cohort (N = 314) and the Stanford TNBC cohort (N = 142). For the Spain TNBC cohort, primary tumor samples were collected from the following medical centers: Hospital de Fuenlabrada - CNIO, Hospital MD Anderson Madrid - CNIO, Hospital Nacional Guillermo Almenara Irigoyen - CNIO, Hospital La Princesa - CNIO, and Hospital 12 de Octubre - CNIO. Samples were deposited and accessed from the CNIO biobank. For the Stanford TNBC cohort, primary tumor samples were collected from the Stanford hospital. Archival formalin-fixed paraffin-embedded tissue samples from TNBC patients (ER and PR positivity less than 1%, HER2 unamplified) were manually reviewed by expert breast cancer pathologists, and representative 1.5mm cores were assembled into 17 tissue microarrays for hematoxylin and eosin (H&E) and MIBI-TOF staining. All patients were followed up after surgery or treatment. Clinical prognostic information, including age, tumor grade, TNM, treatment, and time to relapse is summarized in Supplementary Tables 3-4. Relapse-free survival was calculated as the time from diagnosis to the first metastasic recurrence. Patients without events were censored from the time point of last follow-up. An end date of 1/1/2016 was used for capping when necessary. All of the subsequent staining, preprocessing, and analysis was performed on both the Spain and Stanford TNBC cohorts.

#### Control tissue samples

To assess slide-to-slide variation in staining, we included thirteen 1.5mm control tissue cores on each tissue microarray. Control tissues were carefully selected from the Stanford Pathology Department and included two replicates of tonsil, spleen, lymph node, breast, colon, placenta, and an additional tonsil for asymmetry. Tissue microarrays containing both TNBC samples and control tissues went through all processing steps simultaneously.

#### MIBI-TOF staining and acquisition

##### Panel construction

All antibodies used in this study were previously validated for MIBI-TOF imaging [51, 15, 12]. Antibodies were metal-labeled using the Ionpath conjugation kit (IonPath, Menlo Park, USA) following the manufacturer instructions. To reduce reagent degradation and prolong shelf life, labeled antibodies were then lyophilized individually with 100 mM trehalose in aliquots of 1 *µ*g or 5 *µ*g. Following lyophilization, the appropriate antibody titer was determined by either (1) serial dilution for the new targets (1 *µ*g/mL, 0.5 *µ*g/mL, 0.25 *µ*g/mL, 0.125 *µ*g/mL) or (2) the recommended titer for the relabeled MIBI-TOF validated antibodies. For more details regarding the antibodies, reporter isotopes, or concentrations used in this study, see Supplementary Table 2.

##### Cohort staining

Cohort staining was performed as previously described in Ref. [80]. Briefly, to reduce the potential for batch effects, fresh aliquots of each antibody were reconstituted and combined together into a single mastermix. This mastermix was then used to stain all tissue microarrays. Each step of the staining protocol was performed in pairs, with one reader and one pipettor to reduce mistakes. For more details on the experimental methodology, see the following interactive protocols for Reagent preparation, IHC staining, MIBI staining, and Antibody lyophilization.

##### MIBI-TOF data acquisition

Quantitative MIBI-TOF imaging was performed using a commercial MIBIScope instrument (IonPath, Menlo Park, USA) with 8.0v beam current and 0.63ms dwell time to balance acquisition speed with image clarity. Cores on each tissue microarray were randomly imaged to mitigate potential batch effects due to instrument drift. When possible, images were acquired at 800 *µ*m^2^ at 2048 × 2048 pixels; however, if insufficient tissue was present than an image was acquired at 400 *µ*m^2^ at 1024 × 1024 pixels.

#### Image preprocessing

##### Image compensation and noise removal

MIBI-TOF data often contain several sources of background noise, including gold ions from the conductive slide, organic hydrocarbons, isotropic impurities, and elemental contamination. To systematically correct for any sources of contamination, we performed Rosetta image compensation using the toffy v0.2.2 package in Python [12]. Briefly, coefficients for sources of noise spillover were empirically determined using a gridsearch and Spearman rank correlation and multipliers were chosen and applied to those coefficients by selecting the magnitude that would remove noise while still retaining the actual signal. Each target channel in each image was denoised by subtracting out aforementioned contaminates (Supplementary Figure 8). For precise details on image compensation, see https://github.com/angelolab/publications/tree/main/2024-Ranek_etal_QUICHE.

##### Median pulse height normalization

Following image compensation, we performed median pulse height normalization to correct for any technical differences in MIBI-TOF ion detector sensitivity. A tuning curve was generated using a PMMA sample with fixed ratios of metal isotopes to ensure that any change in signal over the run was due purely to changes in the instrument setup. We then fit a polynomial to this tuning curve to determine the fraction of maximum sensitivity at which the instrument was operating when each channel in each image was acquired. For normalization, we computed the median pulse height for each channel in each image during the run and entered these values into the sensitivity tuning curve to determine the per-mass, per-image sensitivity. This sensitivity estimate was then used to normalize each image (Supplementary Figure 9a-b). Median pulse height normalization was performed using the toffy v0.2.2 package in Python.

##### Quality control

To identify potential sources of batch effects, we measured the slide-to-slide variation in signal intensity across control tissues by calculating the change in the average normalized non-zero pixel intensity for each channel. Overall, we observed minor differences in the signal intensity in control tissues across slides (Supplementary Figure 9c). Using the same metric, we also detected limited spatial bias in the signal intensity across cores on the same slide (Supplementary Figure 9d).

#### Cellular segmentation

Mesmer [53] is a deep learning cell segmentation algorithm that uses a convolutional neural network trained on nuclear (i.e. H3K9ac, H3K27me3) and membrane (i.e. E-cadherin, CD14, CD38, CD45, CK17) images to robustly segment whole cells in multiplexed images of tissues. By incorporating nuclear and cell membrane markers in segmentation, Mesmer can precisely delineate the shape of the cytoplasm of each cell in the tissue. We used Mesmer to identify and segment cells with different morphologies, including epithelial, immune, and stromal cell types (Supplementary Figure 10). Mesmer was implemented using ark-analysis v0.7.0 package in Python under default parameters.

#### Cellular phenotyping

Pixie [81] is an unsupervised pixel clustering algorithm that uses the subcellular localization of proteins to improve cell classification from multiplexed image data. We used pixie to phenotype cells in a four-step process. First, pixels were over-clustered in each image by training a self-organizing map [82] (17 × 17 grid or 289 clusters) on the phenotypic lineage marker pixels across all images in the cohort, including CD45, SMA, Vim, FAP, Fibronectin, Collagen1, CK17, ECAD, ChyTr, Calprotectin, CD3, CD4, CD8, CD11c, CD14, CD20, CD31, CD56, CD63, CD163, HLADR, and FOXP3. These pixel clusters were then grouped into 33 pixel metaclusters using consensus hierarchical clustering [83]. Next, for each cell in the image, we counted the proportion of pixels in each pixel metacluster to generate a *cell* × *pixel metacluster* matrix. Following the same procedure, cells were then over-clustered by training a self-organizing map (17 × 17 grid or 289 clusters) on this pixel metacluster matrix, then grouped into twenty cell metaclusters using consensus hierarchical clustering. In this study, we used the pre-trained self-organizing map weights generated from a large-scale TNBC dataset [52] profiled with the same antibody panel to initialize our clustering results. We validated these predictions through manual inspection of cell type-to-marker expression using Mantis Viewer v1.2.4alpha.3 [84]. Pixie was implemented using the ark-analysis v0.7.0 package in Python. For more details on cell classification, see https://github.com/angelolab/publications/tree/main/2024-Ranek_etal_QUICHE.

#### NeoTRIP TNBC cohort data preprocessing

As a secondary validation, we analyzed 119 baseline tumor samples from the NeoTRIP chemotherapy trial [60, 61] (Supplementary Figure 12). Publicly available imaging mass cytometry and clinical response data (i.e. complete pathologic response (pCR), residual disease (RD)) were accessed from the Zenodo data repository https://doi.org/10.5281/zenodo.7990870. For downstream comparisons to the Spain and Stanford TNBC cohorts, TME and invasive cell types were grouped into higher-order categories as follows. APC: PD-L1^+^APCs, PD-L1^+^IDO^+^APCs. B: CD20^+^B. CD4T: CD4^+^TCF1^+^T, CD4^+^PD1^+^T. CD8T: CD8^+^T, CD8^+^PD1^+^T_*Ex*_, CD8^+^GZMB^+^T, CD8^+^TCF1^+^T. CD163 Mac: M2 Mac. Cancer 4: CD56^+^NE, CD15^+^, Basal, PD-L1^+^IDO^+^, TCF1^+^, PD-L1^+^GZMB^+^, CA9^+^Hypoxia, CA9^+^, Vimentin^+^EMT, CK8/18^*med*^, CK^*lo*^GATA3^+^, pH2AX^+^DSB, Apoptosis, panCK^*med*^, Helios^+^, CK^*hi*^GATA3^+^, MHC I&II^*hi*^, AR^+^LAR. DC: DC. Endothelium: Endothelial. Fibroblast: Myofibroblasts, Fibroblasts. NK: CD56^+^NK. Neutrophil: Neutrophil. PDPN: PDPN^+^Stromal. Plasma: CD79a^+^Plasma. Treg: Treg.

#### Differential cell type abundance analysis

To assess if there were differences in abundance of cell types across patient groups (e.g. relapse/no relapse, pCR/RD), we conducted three complementary analyses. First, we compared the proportions of cell types in patient samples across patient groups in each TNBC cohort using a two-sided Wilcoxon rank sum test (Supplementary Figure 13a, d, g). *p*-values were adjusted using Benjamini-Hochberg false discovery rate correction [85] implemented in the statsmodels v0.14.4 package in Python. Next, we performed differential abundance testing with Milo. Milo [86] is a statistical differential abundance testing method that uses graph neighborhoods to identify cell states that are differentially-enriched across experimental conditions. We implemented Milo using the Pertpy v0.6.0 package in Python [87] to test if there were differences in the abundance of cell types across patient groups in the Spain, Stanford, and NeoTRIP TNBC cohorts. For each cohort, we constructed a *k*-nearest neighbor graph (*k* = 50) of cells according to pairwise Euclidean distances between cells based on their first 20 principal components according to the expression of phenotypic markers. We then annotated cell type neighborhoods by the most abundant cell type (Supplementary Figure 13b, e, h). Lastly, we tested whether the proportion of specific cell types within primary tumor samples were predictive of a patient’s treatment response (NeoTRIP) or recurrence status (Spain and Stanford). For each TNBC cohort, we trained a logistic regression classifier [70] on the proportion of cell types in each patient sample and compared the accuracy of predictions to the ground truth outcome labels by computing the area under the receiver operator curve (AUC). Here, we performed nested 5-fold cross validation to obtain a distribution of predictions. This was repeated over 5 random trials (Supplementary Figure 13c, f, i). Logistic regression classification was implemented using the glmnet v4.1-8 package in R 4.4.0.

#### Immune infiltration analysis

To analyze the coordination of immune cell infiltration within the TNBC primary tumor microenvironment, we analyzed the co-occurrence of immune cell type pairs (e.g. Mono Mac – CD8T, CD8T – CD4T, CD4T – B, B – NK) in primary tumor samples from all three cohorts. Immune cell populations were classified as present or absent in each tumor sample across a range of positivity thresholds (10, 30, 50 cells). A Chi-square test was then used to determine if there were statistically significant differences in immune infiltration across cell type pairs using the scipy v1.12.0 package in Python.

### TNBC QUICHE analysis

#### Differentially-enriched cellular niches at the tumor-immune border

To determine cellular interactions that are differentially-enriched at the invasive margin, we performed differential spatial enrichment analysis using QUICHE as follows. First, we defined structural regions in each tumor (i.e. cancer core, cancer boundary, stroma) by taking the union of the E-cadherin (ECAD) signal with the cancer cell segmentation masks. Tumor compartment masks were then thresholded (0.0015), binarized, and eroded by 100 pixels to define cancer cells within the core and border. All of the remaining cells were defined as belonging in the stroma. Next, we tested for differences in abundance of local cellular niches at the tumor-immune border as compared to the stroma using QUICHE (n_neighbors = 30, radius = 200, k_sim = 100, sketch_size = 1000). QUICHE niche neighborhoods were considered significant if spatial FDR < 0.05. Image processing was performed using the scikit-image v0.19.3 package in Python.

#### Differentially-enriched cellular niches in the extracellular matrix

To determine cellular niches differentially-enriched in aligned/unaligned extracellular matrix (ECM) regions, we performed spatial differential enrichment analysis using a two-step process. First, Collagen1 fibers were segmented from each image using our fiber segmentation algorithm in ark-analysis v0.7.0 in Python [15]. Briefly, Collagen1 images were preprocessed by applying (1) a Gaussian blur (*σ* = 2) to reduce noise, (2) local contrast adjustment (CSD = 256) to account for any differences in fiber signal intensity across the image, and (3) Frangi filtering [88] to enhance fiber-like structures (fiber_widths = (1, 3, 5, 7, 9), min_fiber_size = 100, ridge_cutoff = 0.1). Next, Collagen1 fiber objects were segmented from preprocessed Collagen1 images using the watershed segmentation algorithm [89, 90]. More specifically, dilated and eroded versions of each preprocessed image were created and subjected to multi-Otsu thresholding [91] to classify intensity regions. An elevation map was then generated using the Sobel gradient [92] (Sobel blur = 1) of the smoothed and preprocessed images to highlight edges and enable accurate segmentation of Collagen1 fiber objects with the watershed algorithm. Once fiber objects were extracted and segmented, their length, global orientation, perimeter, and width were computed for each object using `regionprops_table` function in scikit-image v0.19.2. We measured fiber-to-fiber alignment for each fiber in an image, *f*, by calculating the squared distance between the fiber’s orientation *o*_*f*_ and its *k*-nearest fibers as,

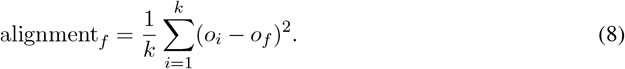

In our analyses, a small alignment distance (alignment_*f*_ ≤ 0.5) indicated that fiber *f* was locally aligned with its (*k* = 4) nearest neighbors. Lastly, we tested for differences the abundance of local cellular niches in aligned or unaligned fiber regions using QUICHE (n_neighbors = 10, radius = 50, k_sim = 50, sketch_size = 500). QUICHE niche neighborhoods were considered significant if spatial FDR < 0.05.

#### Differentially-enriched cellular niches associated with patient outcomes

We used QUICHE to determine cellular interactions that were differentially-enriched in patients that did/did not relapse (Spain and Stanford) or did/did not respond to chemotherapy as measured by complete pathologic response (NeoTRIP). For each cohort, QUICHE was implemented using the following parameters (n_neighbors = 30, radius = 200, k_sim = 100, sketch_size = 1000). Given that the Spain TNBC cohort is a multi-center cohort, we included study center as well as recurrence status as covariates in the negative binomial generalized linear model. QUICHE niche neighborhoods were considered significant if spatial FDR < 0.05. Of note, predicted outcome-associated niche neighborhoods were not correlated with the number of cells within each tumor sample (Supplementary Figure 18a). Moreover, permuted outcome labels achieved a log2 fold change score centered around zero and average spatial FDR corrected *p*-value > 0.05 across 20 random trials highlighting QUICHE’s robustness to false positives (Supplementary Figure 18c). Lastly, to assess the representative accuracy of QUICHE niches in the Spain TNBC cohort, we computed the ratio of recurrence-associated to recurrence-free niches within each patient sample over a range of abundance thresholds. Patients that had more recurrence-associated than recurrence-free niches received a predicted relapse label of 1. We then compared the accuracy of predictions to the ground truth annotations to determine the number of patients that could be classified by QUICHE niche neighborhoods (Figure 5m).

#### Cox proportional hazards model

To assess the generalizability and predictive accuracy of QUICHE niche neighborhoods, we trained a Cox proportional hazards model [93] on the abundance of recurrence-associated QUICHE niches in the Spain TNBC cohort. We then used this trained model to predict recurrence risk for each patient in the Stanford TNBC cohort. Patients were subsequently stratified into high- or low-risk groups based on the median risk score, and a log-rank test was used to test for significant differences in relapse-free survival between risk groups. Kaplan-Meier survival curves [94] and Cox proportional hazards regression models were generated using the survival v3.5-8 package in R 4.4.0.

## Data Availability

MIBI-TOF data and cell segmentation masks for both the Spain and Stanford TNBC cohorts are publicly available in the BioStudies repository under the accession number S-BIAD1507 at https://www.ebi.ac.uk/biostudies/bioimages/studies/S-BIAD1507 [95]. Preprocessed anndata objects for all three TNBC cohorts are publicly available in the Zenodo repository at https://doi.org/10.5281/zenodo.14290163 [96].

## Code Availability

QUICHE is implemented as an open-source Python package and is publicly available at https://github.com/jranek/quiche. Source code including all functions for benchmarking evaluation and figure generation are publicly available at https://github.com/angelolab/publications/tree/main/2024-Ranek_etal_QUICHE.

## Acknowledgments

The authors would like to thank all patients and study personnel. The authors thank Erin McCaffrey, Seongyuel Park, and Kiyan Abel for their insightful discussions related to this work. This work was supported by NIH grants 5U54CA20997105 (M.A.), 5DP5OD01982205 (M.A.), 1R01CA24063801A1 (M.A.), 5R01AG06827902 (M.A.), 5UH3CA24663303 (M.A.), 5R01CA22952904 (M.A.), 1U24CA22430901 (M.A.), 5R01AG05791504 (M.A), 5R01AG05628705 (M.A.); the Department of Defense W81XWH2110143 (M.A.), the Wellcome Trust (M.A.) and other funding from the Bill and Melinda Gates Foundation (M.A.), Cancer Research Institute (M.A.), the Parker Center for Cancer Immunotherapy (M.A.), and the Breast Cancer Research Foundation (M.A.).

## Author Contributions

J.S.R and M.A. conceptualized this study. S.M. and M.Q.F. provided samples and clinical data for the Spain TNBC cohort. R.B.W. provided samples and clinical data for the Stanford cohort. N.F.G, C.C., and M.G. generated the MIBI-TOF data. J.S.R performed the data preprocessing, benchmarking, evaluation, analysis, and wrote the manuscript. C.S. developed the fiber segmentation algorithm. A.K. performed Rosetta image compensation. M.A. supervised the project. All authors read and provided feedback on the manuscript.

