## Supplementary Information for "QUICHE reveals structural definitions of anti-tumor responses in triple negative breast cancer"

### Supplementary Tables

**Supplementary Table S1: Guidelines on QUICHE parameter selection**

| Parameter | Description | Required | Recommendation |
|---|---|---|---|
| | | *Niche Identification* | |
| n_neighbors | Encodes density and spatial proximity in niche determination | N (recommended) | 30 neighbors or 3-hop |
| radius | Bounds the local composition of niches | Y | 100-300 pixels |
| sketch size | Number of niches from each sample for condition-specific comparisons | Y | L1 elbow |
| | | *Graph Construction and Differential Enrichment Analysis* | |
| $k$_sim | Defines neighborhood size in niche similarity graph | Y | Data-dependent (50-300) |
| model contrasts | Specifies condition-specific comparisons for differential niche analysis | Y | Analysis-dependent |
| | | *QUICHE Niche Neighborhood Annotation* | |
| n_largest | Labels niche neighborhoods according to most abundant cell types | N (recommended) | Top 3 |

#### Supplementary Table S2: Antibody panel information

| Internal ID | Target | Clone | Vendor | Cat Number | Lot | Mass | Element | Titer ($\mu$g/mL) |
|---|---|---|---|---|---|---|---|---|
| 1352 | Calprotectin | MAC387 | Thermofisher | MA1-80446 | VC2962354 | 69 | Ga | 2 |
| 1350 | Mast Cell Chymase | EPR13136 | Abcam | ab233729 | GR32553931 | 71 | Ga | 1 |
| 1351 | Mast Cell Tryptase | EPR9522 | Abcam | ab181724 | GR32471283 | 71 | Ga | 1 |
| 1701 | SMA | D4K9N | Cell Signaling Technology | 19245BF | 4 | 89 | Y | 1 |
| 1163 | Vimentin | D21H3 | Cell Signaling Technology | 5741BF | 7 | 113 | In | 2 |
| 1729 | CD4 | EPR6855 | Abcam | ab181724 | GR3285644-16 | 143 | Nd | 0.5 |
| 1784 | CD69 | EPR21814 | Abcam | ab234512 | GR3375521 | 144 | Nd | 0.25 |
| 1603 | FAP | poly | R&D Systems | AF3715 | zkw0619101 | 145 | Nd | 0.5 |
| 1567 | FOXP3 | 236AE7 | Abcam | ab96048 | GR3215102-3 | 146 | Nd | 1 |
| 1773 | PD1 | D4W2J | Cell Signaling Technology | 86163BF | 7 | 147 | Sm | 1 |
| 1564 | CD31 | EP3095 | Abcam | ab226157 | GR3356121-6 | 148 | Nd | 0.25 |
| 1362 | Biotin | 1D4-C5 | Biolegend | 409002 | B267484 | 149 | Sm | 1 |
| 1272 | Ecadherin | 24E10 | Cell Signaling Technology | 3195BF | 16 | 150 | Nd | 0.25 |
| 1514 | CD56 151Eu | MRQ42 | Ionpath | 715101-100 | 19235-02(1) | 151 | Eu | 0.5 |
| 1721 | CD38 | E7Z8C | Cell Signaling Technology | 51000BF | 2 | 152 | Sm | 0.05 |
| 1730 | TCF1 TCF7 | C63D9 | Cell Signaling Technology | 2203BF | 12 | 153 | Eu | 0.25 |
| 1665 | TBET | D6N8B | Cell Signaling Technology | 13232BF | 6 | 154 | Sm | 0.5 |
| 1783 | CD45RB | MEM55 | Biolegend | 310202 | 328313 | 155 | Gd | 0.5 |
| 1164 | CD68 | D4B9C | Cell Signaling Technology | 76437BF | 2 | 156 | Gd | 0.5 |
| 1754 | CD11c | EP1347Y | Abcam | ab216655 | GR3357092-12 | 157 | Gd | 1 |
| 1462 | CD8 | C8144B | Cell Marque | 108M-OEM1404 | | 158 | Gd | 0.5 |
| 1292 | CD3e | D7A6E | Cell Signaling Technology | 8506BF | 4 | 159 | Tb | 1 |
| 1104 | IDO1 | ab55305 | Abcam | ab55305 | GR239251-1 | 160 | Gd | 0.5 |
| 1310 | CD45RO | UCHL1 | Ionpath | 716101-100 | 19235-05 | 161 | Dy | 0.5 |
| 1593 | TIM-3 | EPR22241 | Ionpath | 716201-100 | 20238-02(3) | 162 | Dy | 0.5 |
| 1295 | CD163 | D6U1J | Cell Signaling Technology | 93498BF | 2 | 163 | Dy | 0.5 |
| 1198 | CD20 | L26 | Cell Marque | 120M-8-OEm | 1908402 | 164 | Dy | 0.25 |
| 1749 | FN1 | E5H6X | Cell Signaling Technology | 26836BF | 2 | 165 | Ho | 2 |
| 1740 | Glut1 | EPR3915 | Abcam | ab252403 | GR3374509-1 | 166 | Er | 0.5 |
| 1285 | HLADR | EPR3692 | Abcam | ab215985 | GR3218615-1 | 167 | Er | 0.25 |
| 1637 | CD14 | D7A2T | Cell Signaling Technology | 56082BF | 2 | 168 | Er | 0.25 |
| 1565 | CD45 | D9M8I | Cell Signaling Technology | 13917BF | 8 | 169 | Tm | 0.25 |
| 1728 | Cytokeratin17 | SP95 | Abcam | ab238808 | GR3286787-3 | 170 | Er | 0.25 |
| 1584 | COL1A1 | E8F4L | Cell Signaling Technology | 72026BF | 2 | 171 | Yb | 1 |
| 1720 | H3K27me3 | C36B11 | Cell Signaling Technology | 9733BF | 20 | 172 | Yb | 1 |
| 1747 | CD57 | NK/804 | Abcam | ab212408 | GR3372900-2 | 173 | Yb | 0.5 |
| 1342 | H3K9ac | C5B11 | Cell Signaling Technology | 9649BF | 12 | 174 | Yb | 1 |
| 1704 | Ki67 | B56 | BD Biosciences | 556003 | 8239549 | 175 | Lu | 0.25 |
| 1723 | HLA1 class ABC | EMR8-5 | Abcam | ab70328 | GR3380465-1 | 176 | Yb | 1 |
| N/A | PDL1 BiotinylatedE1L3N | | Cell Signaling Technology | 15118S | N/A | N/A | N/A | 2 |

**Supplementary Table S3: Spain TNBC cohort characteristics (N = 314)**

| Characteristics | No Relapse (N = 205) | Relapse (N = 109) |
|---|---|---|
| **Age (years)** | | |
| median [min, max] | 54.0 [28.2, 90.5] | 57.7 [24.7, 88.2] |
| **T** | | |
| 1 | 61 (29.8%) | 14 (12.8%) |
| 2 | 114 (55.6%) | 55 (50.5%) |
| 3 | 22 (10.7%) | 18 (16.5%) |
| 4 | 7 (3.4%) | 21 (19.3%) |
| Unknown | 1 (0.5%) | 1 (0.9%) |
| **N** | | |
| 0 | 134 (65.4%) | 39 (35.8%) |
| 1 | 42 (20.5%) | 25 (22.9%) |
| 2 | 21 (10.2%) | 16 (14.7%) |
| 3 | 8 (3.9%) | 23 (21.1%) |
| Unknown | 0 (0.0%) | 6 (5.5%) |
| **Histological grade** | | |
| 1/2 | 4 (1.95%) | 2 (1.83%) |
| 3 | 199 (97.07%) | 106 (97.25%) |
| Unknown | 2 (0.98%) | 1 (0.92%) |
| **Type of chemotherapy** | | |
| Adjuvant | 125 (61.0%) | 54 (49.5%) |
| Neoadjuvant | 58 (28.3%) | 31 (28.4%) |
| No treatment | 22 (10.7%) | 24 (22.0%) |
| **Medical Center** | | |
| Hospital de Fuenlabrada - CNIO | 29 (14.1%) | 11 (10.1%) |
| Hospital MD Anderson Madrid - CNIO | 57 (27.8%) | 33 (30.3%) |
| Hospital Nacional Guillermo Almenara Irigoyen - CNIO | 30 (14.6%) | 16 (14.7%) |
| Hospital La Princesa - CNIO | 13 (6.3%) | 7 (6.4%) |
| Hospital 12 de Octubre - CNIO | 76 (37.1%) | 42 (38.5%) |
| **Time to relapse (days)** | | |
| Median [min, max] | – | 479.5 [33.0, 3839.0] |

**Supplementary Table S4: Stanford TNBC cohort characteristics (N = 142)**

| Characteristics | No Relapse (N = 97) | Relapse (N = 45) |
|---|---|---|
| **Age (years)** | | |
| median [min, max] | 54 [31, 91] | 49 [26, 84] |
| **T** | | |
| 1 | 40 (42.3%) | 13 (28.9%) |
| 2 | 36 (37.1%) | 11 (24.4%) |
| 3 | 2 (2.1%) | 4 (8.9%) |
| 4 | 1 (1.0%) | 1 (2.2%) |
| IS | 1 (1.0%) | 0 (0.0%) |
| X | 17 (17.5%) | 16 (35.6%) |
| **N** | | |
| 0 | 53 (54.6%) | 9 (20.0%) |
| 1 | 17 (18.6%) | 13 (28.9%) |
| 2 | 7 (7.2%) | 1 (2.2%) |
| 3 | 1 (1.0%) | 6 (13.3%) |
| X | 19 (19.6%) | 16 (35.6%) |
| **M** | | |
| 0 | 55 (56.7%) | 18 (40.0%) |
| 1 | 0 (0.0%) | 1 (2.2%) |
| X | 42 (44.3%) | 26 (57.8%) |
| **Stage** | | |
| 1 | 38 (39.2 %) | 4 (8.9 %) |
| 2 | 44 (46.4 %) | 23 (51.1 %) |
| 3 | 13 (13.4 %) | 14 (31.1 %) |
| 4 | 0 (0.0 %) | 4 (8.9 %) |
| Unknown | 2 (2.1 %) | 0 (%) |
| **Grade** | | |
| 1/2 | 21 (21.6 %) | 11 (24.4 %) |
| 3/4 | 71 (74.2 %) | 30 (66.7 %) |
| Unknown | 5 (5.2 %) | 4 (8.9 %) |
| **Time to relapse (days)** | | |
| Median [min, max] | – | 272.0 [0.0, 2507.0] |

### Supplementary Figures

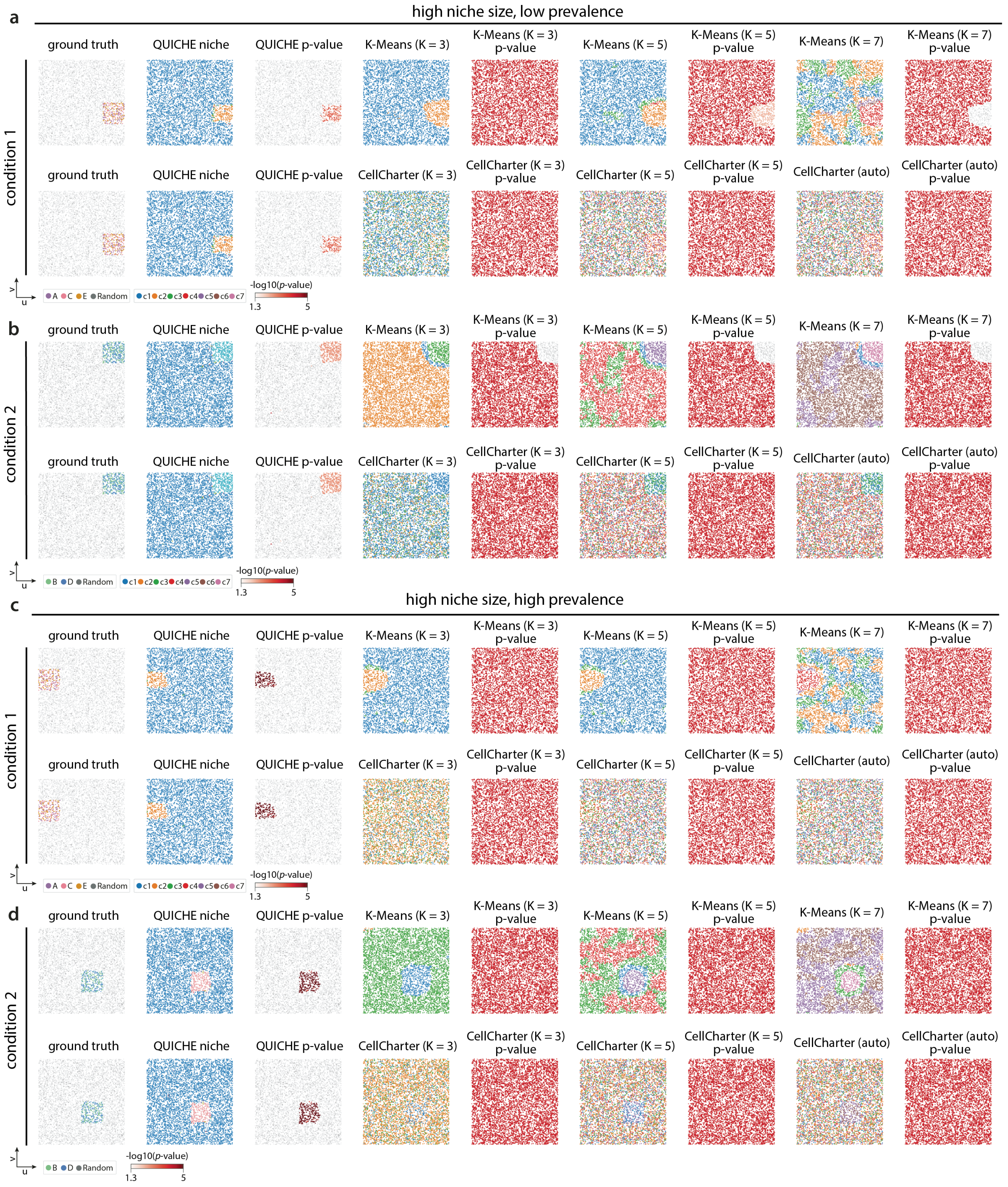


**Supplementary Figure 1: Qualitative comparisons of spatial enrichment methods on unstructured spatial topologies (high niche size/low prevalence, high niche size/high prevalence).** We benchmarked spatial methods on their ability to recover condition-specific cellular niches when varying the underlying spatial topology, niche size, or sample prevalence. (a-b) Performance of different spatial clustering approaches on detecting condition-specific differences in cellular organization on simulated data with unstructured spatial topologies, high niche size (11.1% sample), and low prevalence (40% patient samples). Example visualizations of clustering results and differential enrichment for (a) condition 1 where cell types A, C, and E were differentially-enriched and (b) condition 2 where cell types B and D were differentially-enriched. (c-d) Performance on simulated data with unstructured spatial topologies, high niche size (11.1% sample), and high prevalence (100% patient samples).

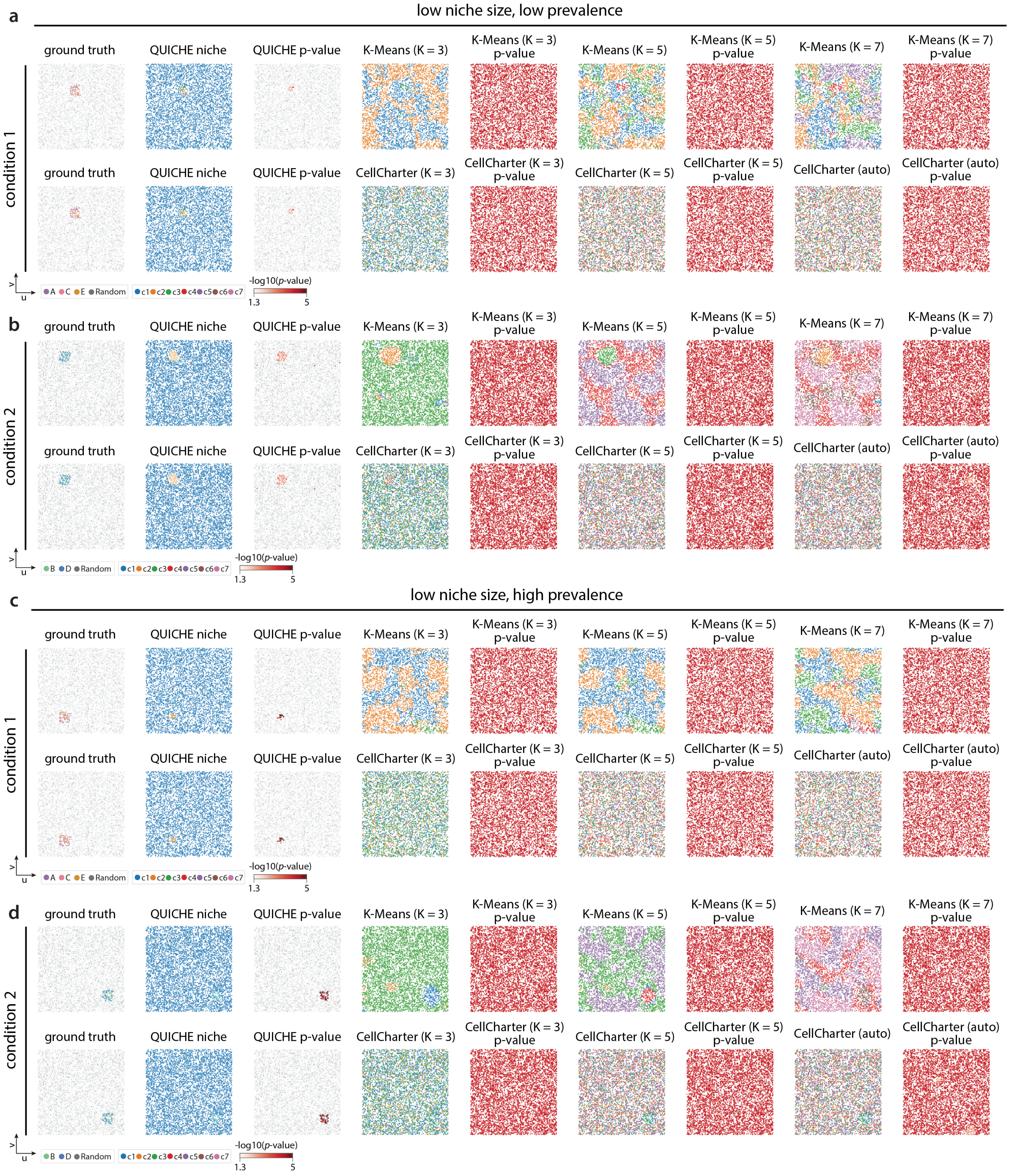


**Supplementary Figure 2: Qualitative comparisons of spatial enrichment methods on unstructured spatial topologies (low niche size/low prevalence, low niche size/high prevalence).** We benchmarked spatial methods on their ability to recover condition-specific cellular niches when varying the underlying spatial topology, niche size, or sample prevalence. (a-b) Performance of different spatial clustering approaches on detecting condition-specific differences in cellular organization on simulated data with unstructured spatial topologies, low niche size (1.6% sample), and low prevalence (40% patient samples). Example visualizations of clustering results and differential enrichment for (a) condition 1 where cell types A, C, and E were differentially-enriched and (b) condition 2 where cell types B and D were differentially-enriched. (c-d) Performance on simulated data with unstructured spatial topologies, low niche size (1.6% sample), and high prevalence (100% patient samples).

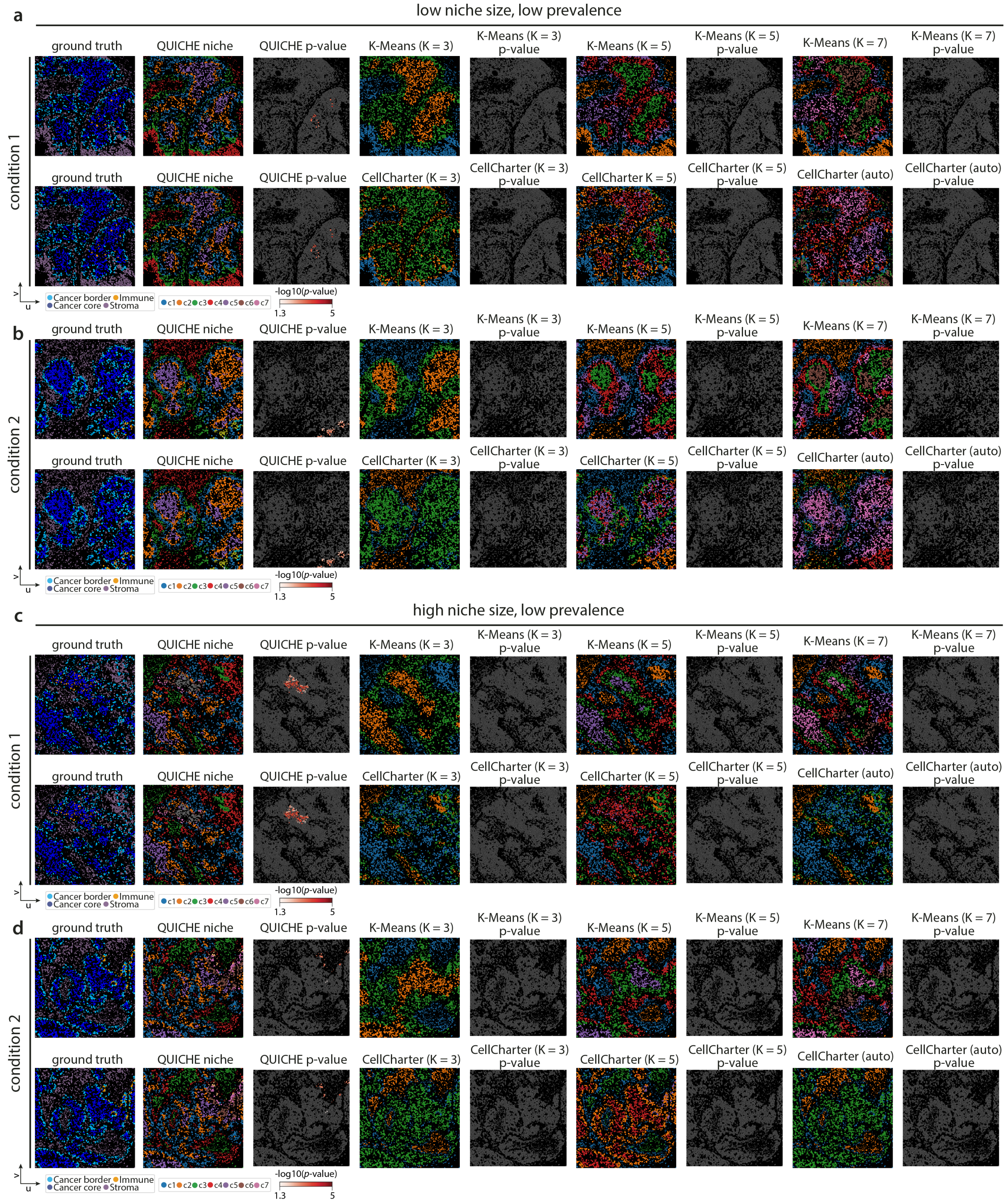


**Supplementary Figure 3: Qualitative comparisons of spatial enrichment methods on structured spatial topologies (low niche size/low prevalence, high niche size/low prevalence).** We benchmarked spatial methods on their ability to recover condition-specific cellular niches when varying the underlying spatial topology, niche size, or sample prevalence. (a-b) Performance of different spatial clustering approaches on detecting condition-specific differences in cellular organization on simulated data with structured spatial topologies, low niche size (25 immune cells within 500 pixel radius), and low prevalence (40% patient samples). Example visualizations of clustering results and differential enrichment for (a) condition 1 where immune cells were differentially-enriched in the cancer core and (b) condition 2 where immune cells were differentially-enriched in the cancer border. (c-d) Performance on simulated data with structured spatial topologies, high niche size (100 immune cells within 500 pixel radius), and low prevalence (40% patient samples).

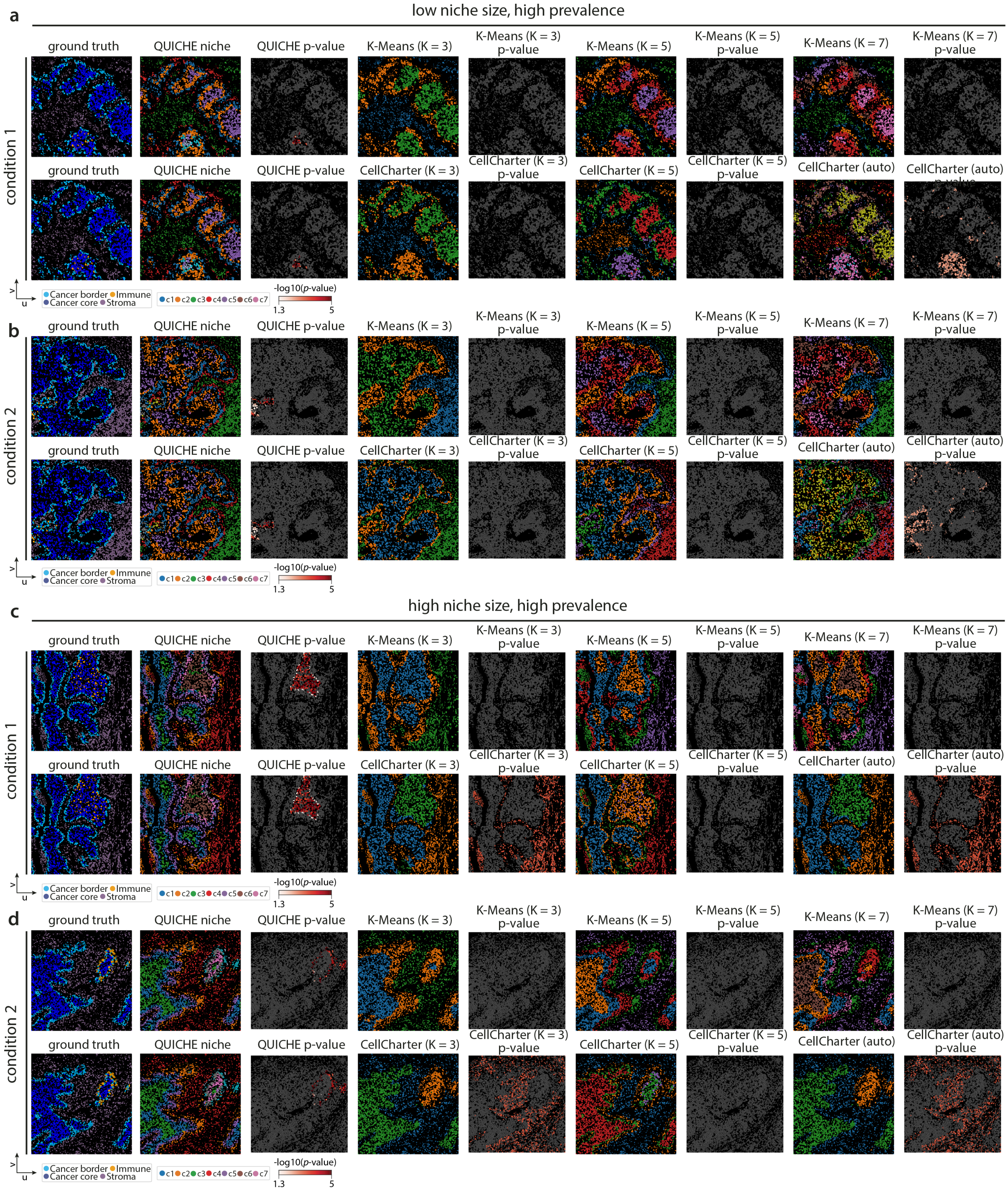


**Supplementary Figure 4: Qualitative comparisons of spatial enrichment methods on structured spatial topologies (low niche size/high prevalence, high niche size/high prevalence).** We benchmarked spatial methods on their ability to recover condition-specific cellular niches when varying the underlying spatial topology, niche size, or sample prevalence. (a-b) Performance of different spatial clustering approaches on detecting condition-specific differences in cellular organization on simulated data with structured spatial topologies, low niche size (25 immune cells within 500 pixel radius), and high prevalence (100% patient samples). Example visualizations of clustering results and differential enrichment for (a) condition 1 where immune cells were differentially-enriched in the cancer core and (b) condition 2 where immune cells were differentially-enriched in the cancer border. (c-d) Performance on simulated data with structured spatial topologies, high niche size (100 immune cells within 500 pixel radius), and high prevalence (100% patient samples).

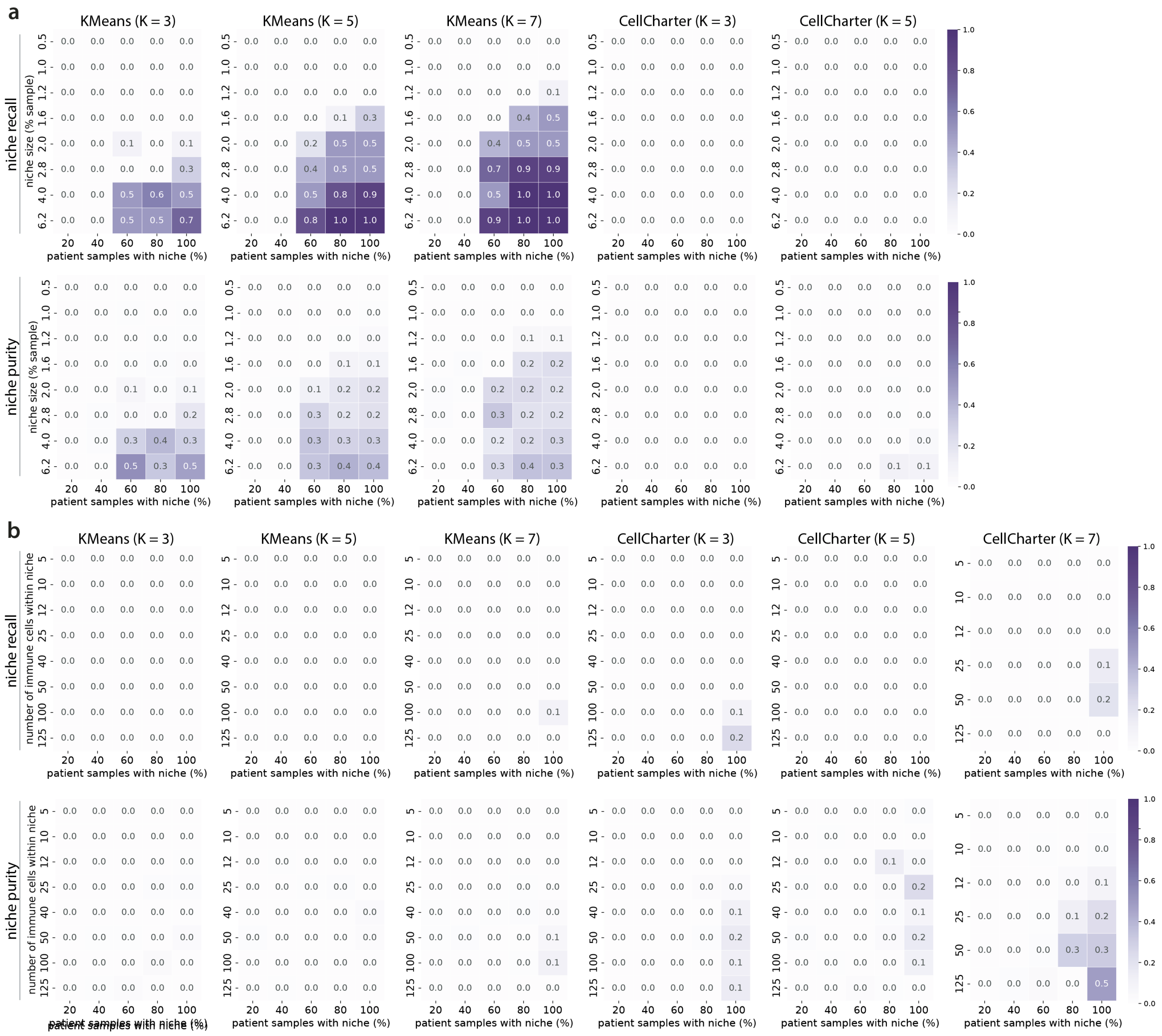


**Supplementary Figure 5: Performance of spatial clustering methods across different clustering resolutions.** Heatmap shows the performance of spatial clustering methods, $K$-Means++ and CellCharter, on detecting differentially-enriched cellular niches on (a) unstructured spatial topologies and (b) structured spatial topologies.

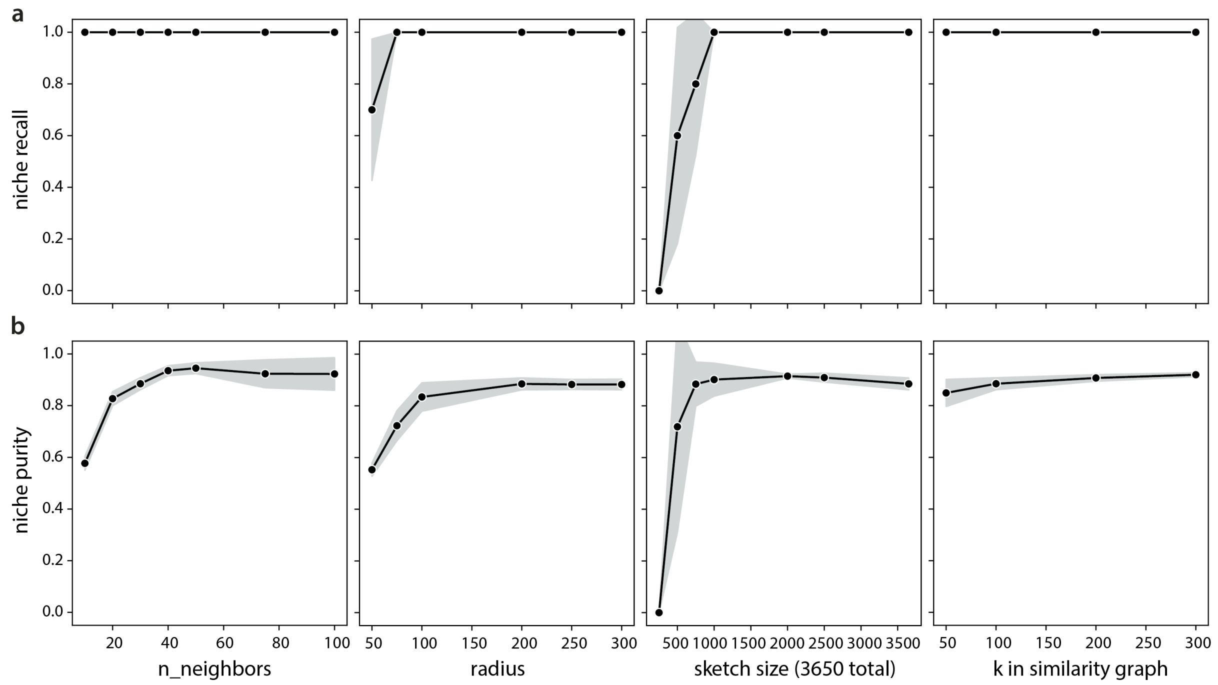


**Supplementary Figure 6: QUICHE is robust to changes in hyperparameters for structured spatial topologies.** Multi-patient spatial proteomic imaging data were simulated with structured spatial topologies with a niche size of 50 immune cells within 500 pixel radius and niche prevalence of 100% samples. QUICHE achieves similar (a) niche recall and (b) niche purity scores across a range of hyperparameters, including the number of nearest neighbors in spatial niche detection, the radius size in spatial niche detection, the number of niches sampled from each patient sample, and the number of $k$-nearest neighbors in niche similarity graph construction. The error bands show the standard deviation over $n = 5$ random trials.

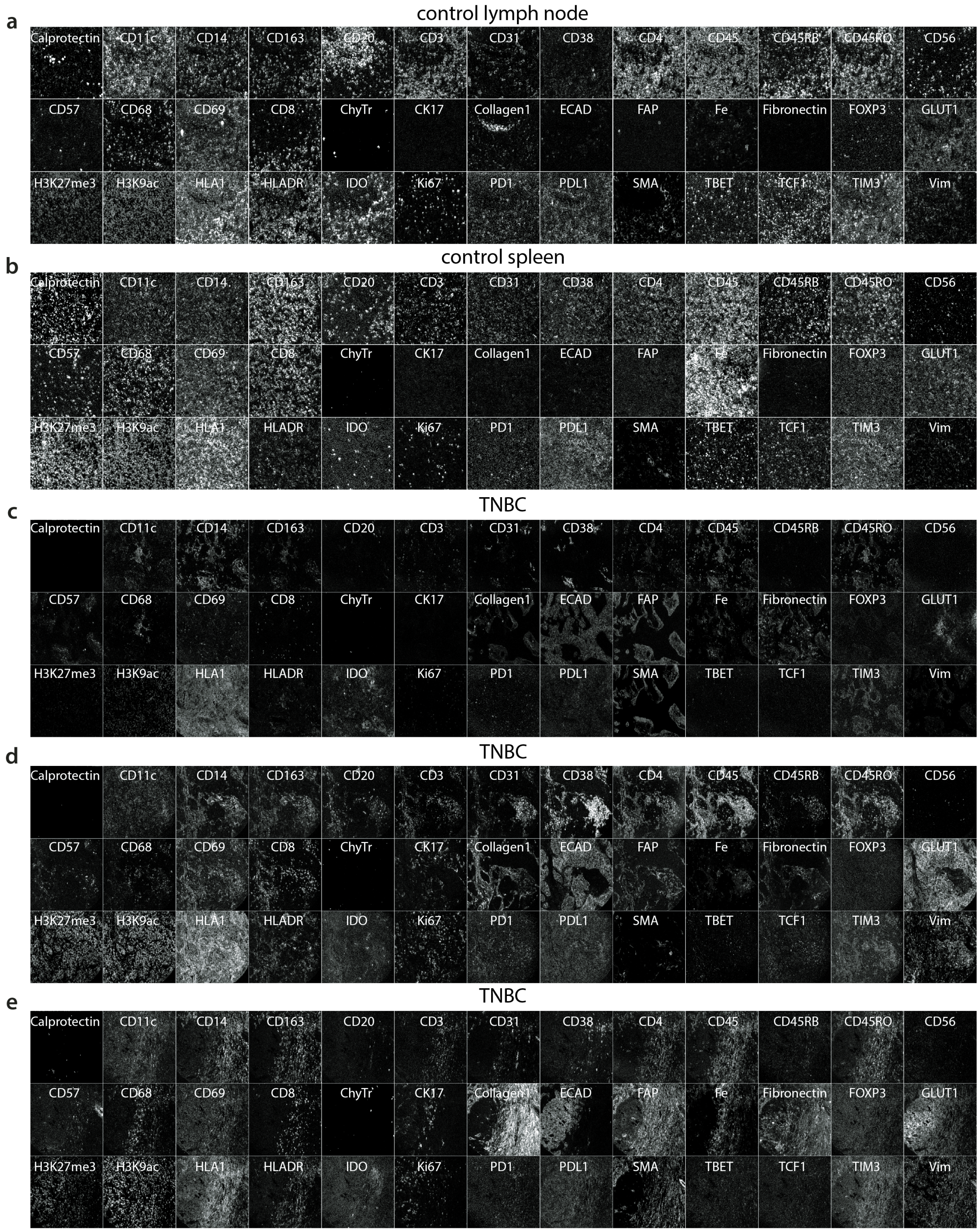


**Supplementary Figure 7: Antibody panel validation.** Single-plex images of profiled proteins in (a) control lymph node sample (b) control spleen sample and (c-e) three representative triple-negative breast cancer samples from the Spain TNBC cohort.

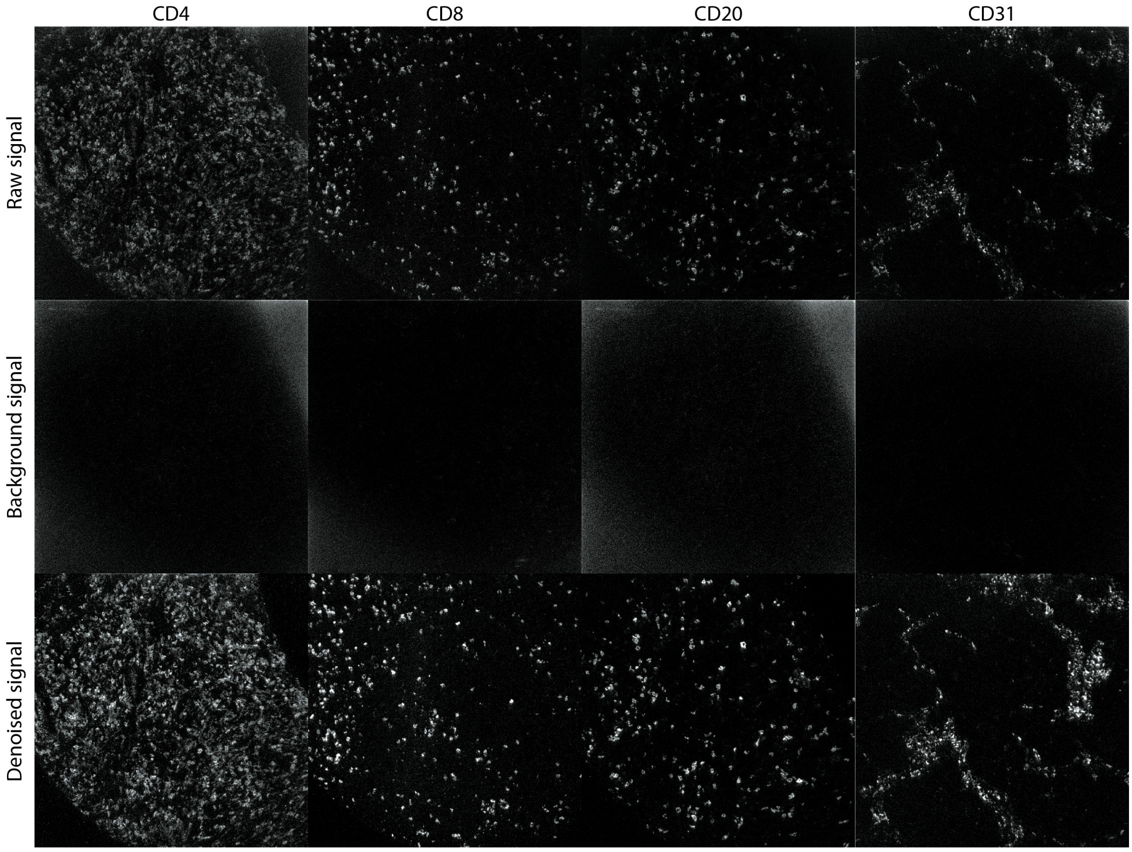


**Supplementary Figure 8: Image compensation with Rosetta.** Representative images of several channels before and after background subtraction with the Rosetta algorithm.

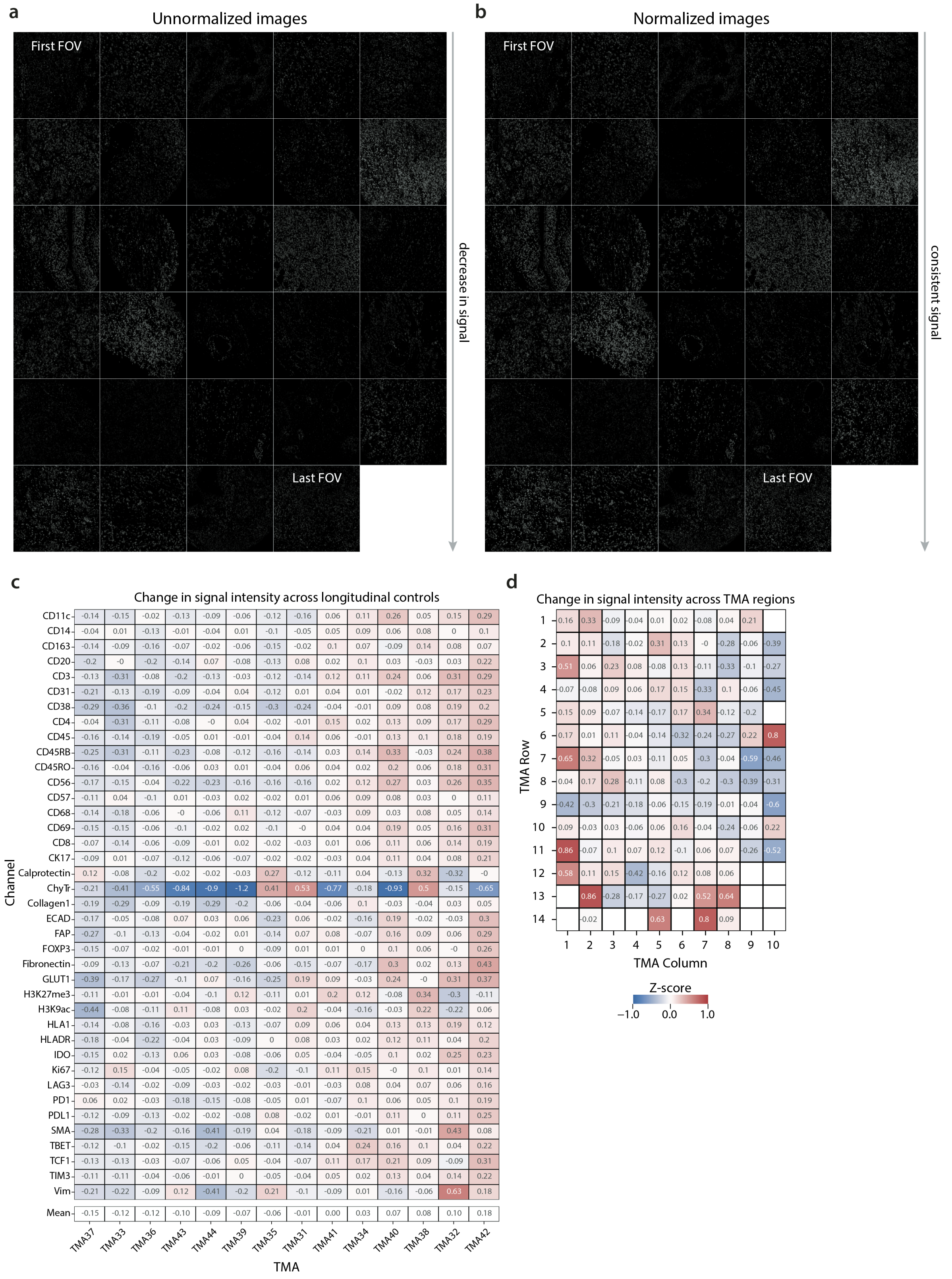


**Supplementary Figure 9: Preprocessing and quality control for the Spain TNBC cohort.** (a) Representative images show the unnormalized intensities from the start of acquisition (top left) to the end of acquisition (bottom right). (b) Representative images show the normalized intensities from the start of acquisition (top left) to the end of acquisition (bottom right) using median pulse height normalization. (c) Heatmap shows the average change in normalized expression of protein markers across control tissues on different tissue microarrays. (d) Heatmap shows the average change in normalized expression across spatial locations of each tissue microarray.

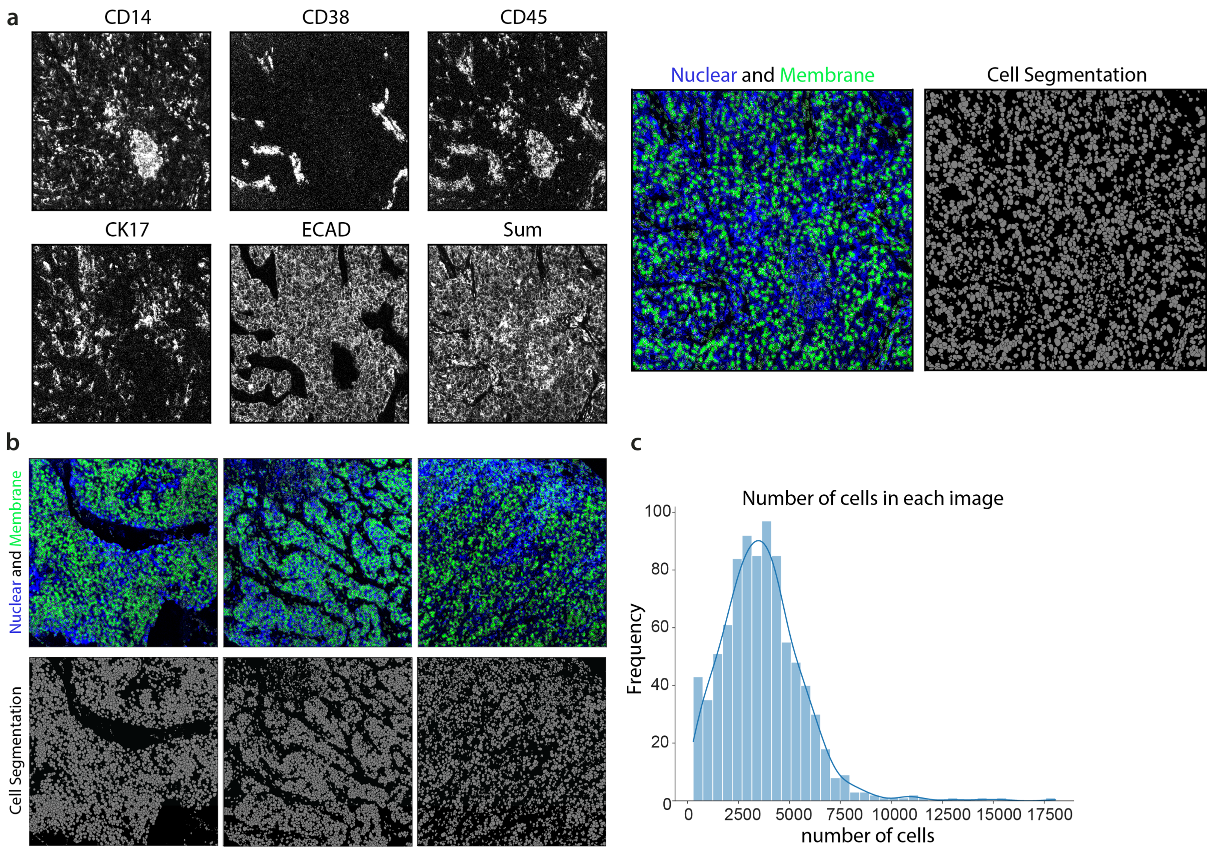


**Supplementary Figure 10: Cell segmentation with Mesmer.** (a) Representative images show the membrane markers (CD14, CD38, CD45, CK17, ECAD) used for cell segmentation, in addition to the nuclear markers (H3K9ac, H3K27me3). (b) Representative images of three tumors annotated by nuclear and membrane markers (top) and predicted cell segmentation masks (bottom). (c) Histogram shows the number of cells in each tumor image in the Spain TNBC cohort.

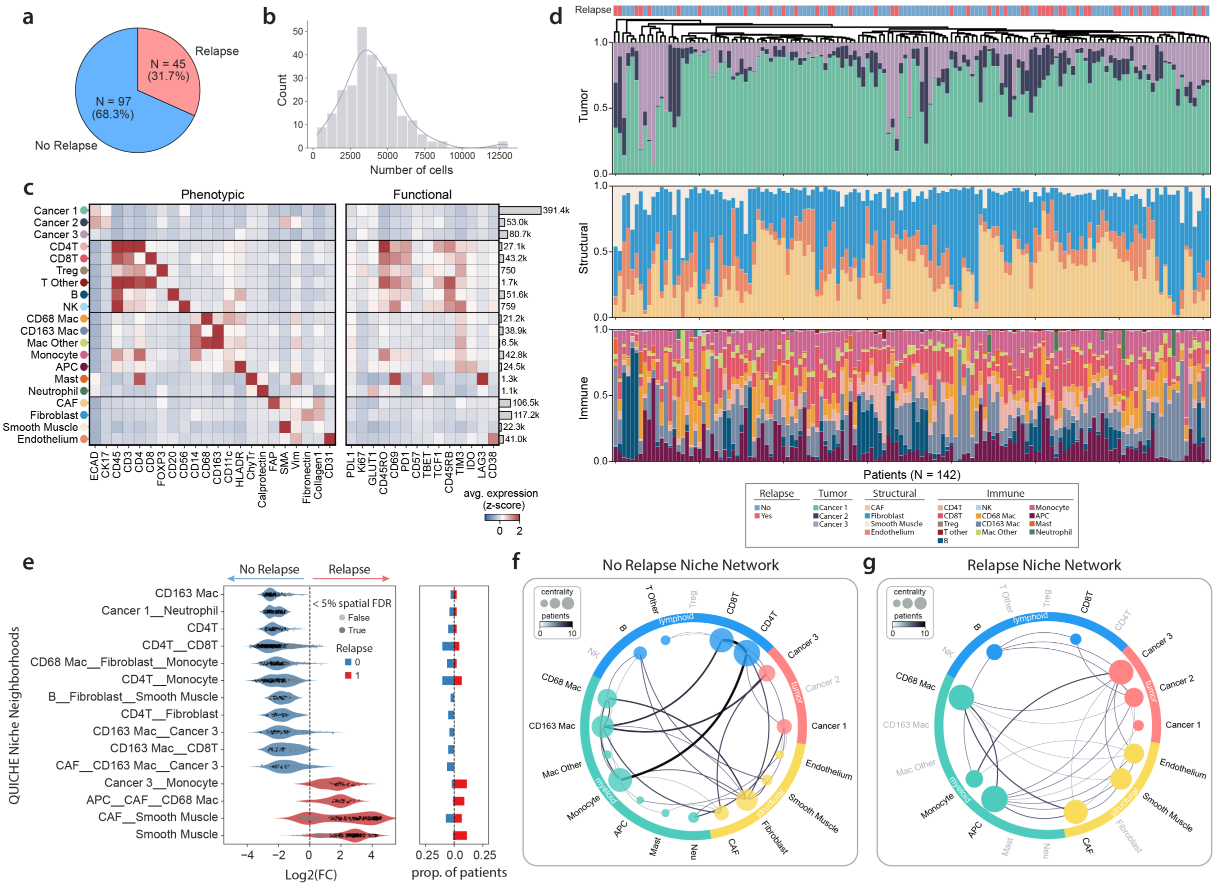

**Supplementary Figure 11: Stanford cohort overview.** (a) Pie chart shows the number of patients that did (pink) or did not relapse (blue). (b) Histogram shows the number of cells in each tumor image. (c) Heatmap shows the average standardized expression of twenty cell phenoytypes clustered according to protein expression of phenotypic markers. Barplots to the right show the total number of cells within each cell phenotype. (d) Barplots show the frequency of tumor (top), structural (middle) and immune lineages (bottom) across all primary tumor samples in this cohort. Samples were hierarchically clustered according all frequency subsets. Relapse status is denoted by color in the top row. (e) Violin plots show the top 15 differentially-enriched niche neighborhoods in patients who relapsed (red) or did not relapse (blue). Barplots to the right show the proportion of patients with a niche neighborhood in the respective patient groups. (f) Niche network for non-relapsing patients, where nodes represent cell types within recurrence-free niches and edge weights correspond to the number of unique patients with the corresponding interaction. Node size is proportional to connectedness, as measured by eigenvector centrality. (g) Niche network for relapsing patients, where nodes represent cell types within recurrence-associated niches and edge weights correspond to the number of unique patients with the corresponding interaction. Node size is proportional to connectedness, as measured by eigenvector centrality.

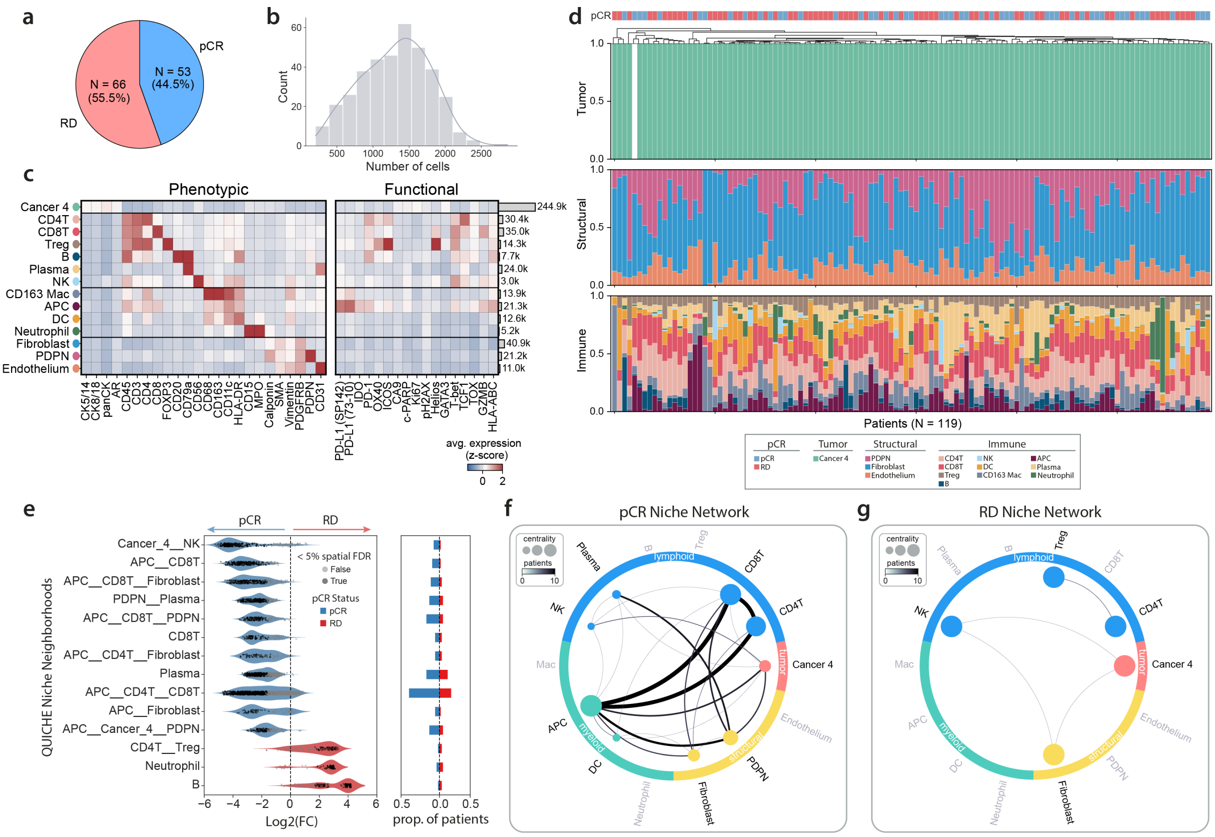


**Supplementary Figure 12: NeoTRIP cohort overview.** (a) Pie chart shows the number of patients that had complete pathologic response (pCR) (blue) or residual disease (RD) (pink). (b) Histogram shows the number of cells in each tumor image. (c) Heatmap shows the average standardized expression of fourteen cell phenoytypes clustered according to protein expression. Cell type annotations from the original study were grouped into higher order categories for downstream comparisons to the Spain and Stanford cohorts. Barplots to the right show the total number of cells within each cell phenotype. (d) Barplots show the frequency of tumor (top), structural (middle) and immune lineages (bottom) across all primary tumor samples in this cohort. Samples were hierarchically clustered according all frequency subsets. pCR status is denoted by color in the top row. (e) Violin plots show the top 14 differentially-enriched niche neighborhoods in patients who had complete pathologic response (blue) or residual disease (red). Barplots to the right show the proportion of patients with a niche neighborhood in the respective patient groups. (f) Niche network for responding patients, where nodes represent cell types within pCR-associated niches and edge weights correspond to the number of unique patients with the corresponding interaction. Node size is proportional to connectedness, as measured by eigenvector centrality. (g) Niche network for patients that had residual disease, where nodes represent cell types within RD-associated niches and edge weights correspond to the number of unique patients with the corresponding interaction. Node size is proportional to connectedness, as measured by eigenvector centrality.

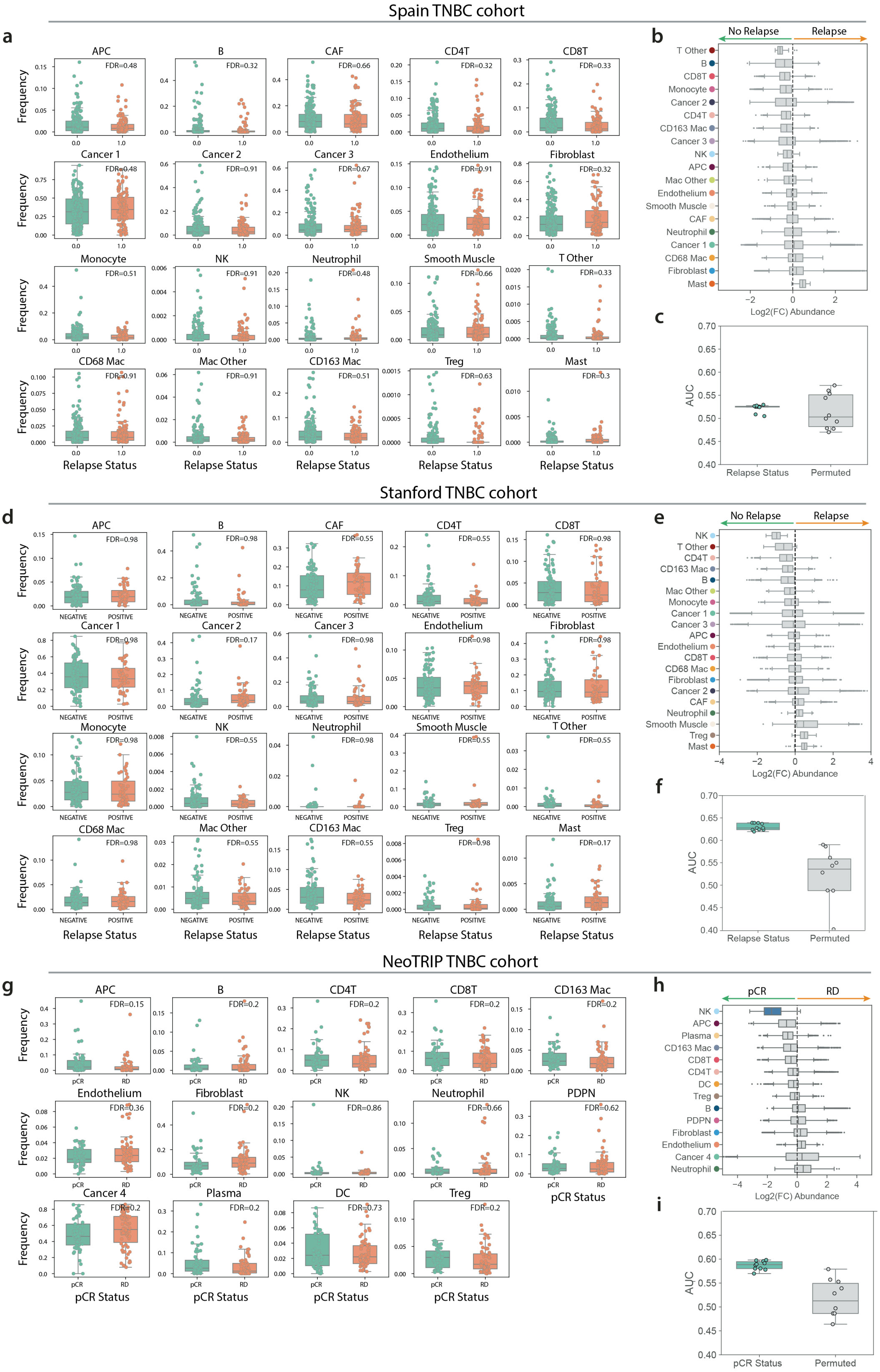


**Supplementary Figure 13: Differential cell type abundance analysis across TNBC cohorts.** (a-c) Differential cell type analysis across recurrence groups in the Spain TNBC cohort. (a) Boxplots show the frequency of cell phenotypes across patients that did (orange) or did not relapse (green). A two-sided Wilcoxon rank sum test was used to test for differences between patient groups. Multiple hypothesis testing correction was performed using Benjamini Hochberg. (b) Differential cell type abundance testing was performed using Milo. Boxplots show the log2 fold change in the abundance of cell type neighborhoods across recurrence groups. (c) A logistic regression classifier was trained on the proportion of cell types to predict relapse status using 5-fold cross validation. Boxplots show the average area under the receiver operator curve (AUC) values across 10 random trials. Permuted label performance is shown in gray. (d-f) Differential cell type analysis across recurrence groups in the Stanford TNBC cohort. (g-i). Differential cell type analysis across pathological complete response (pCR) groups in the NeoTRIP TNBC cohort. RD indicates patients that had residual disease.

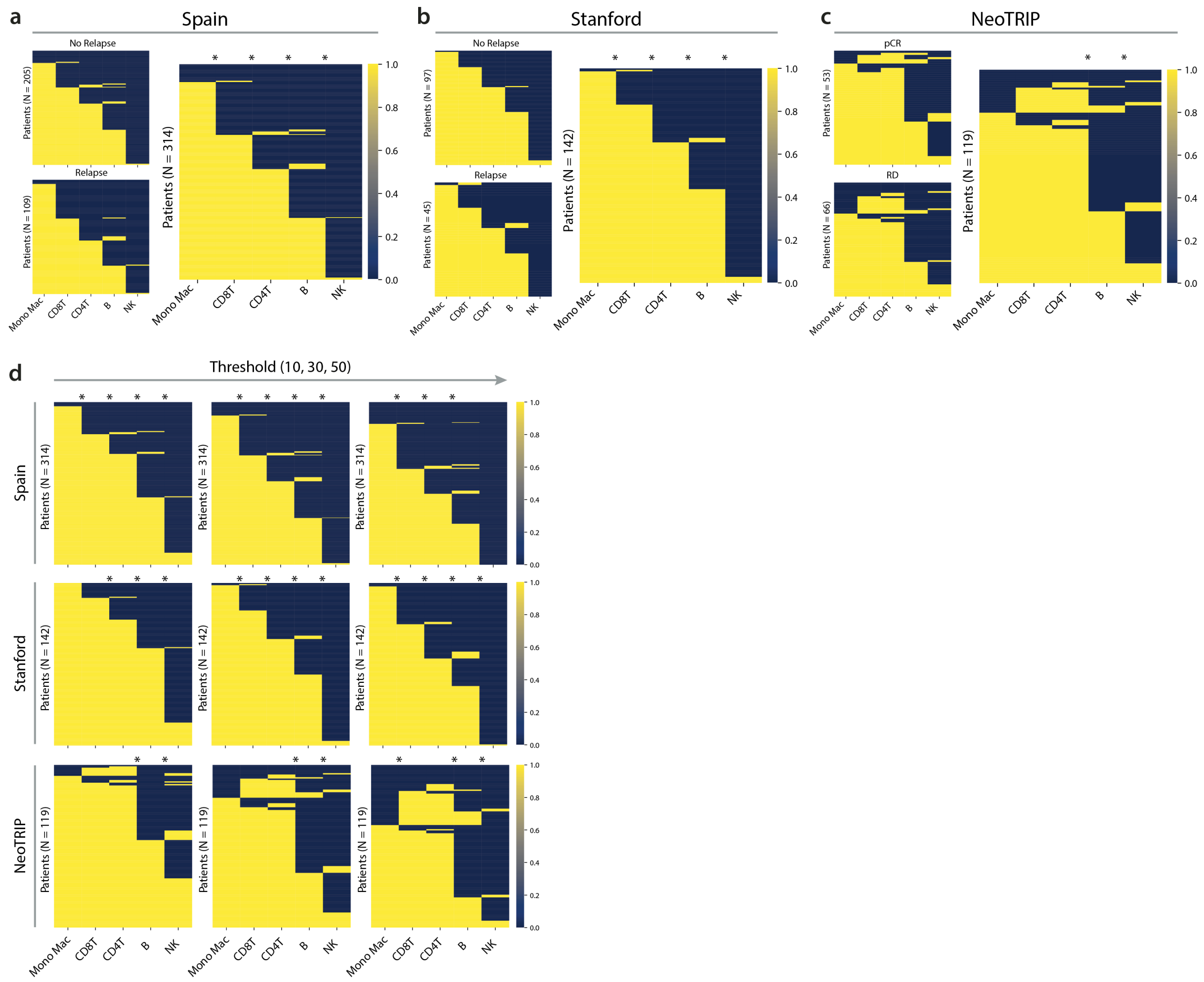


**Supplementary Figure 14: Stepwise immune infiltration across TNBC cohorts.** Heatmaps show the presence (yellow) or absence (blue) of immune cells across patients in the (a) Spain, (b) Stanford, and (c) NeoTRIP TNBC cohorts. For each cohort, a Chi-square test was used to test for differences in the infiltration of cell type pairs (Mono Mac - CD8T, CD8T - CD4T, CD4T - B, B - NK). * indicates $p$-value < 0.05. (d) Heatmap shows immune infiltration with different thresholds for positivity (x-axis) across patients in the Spain (top), Stanford (middle) and NeoTRIP (bottom) TNBC cohorts.

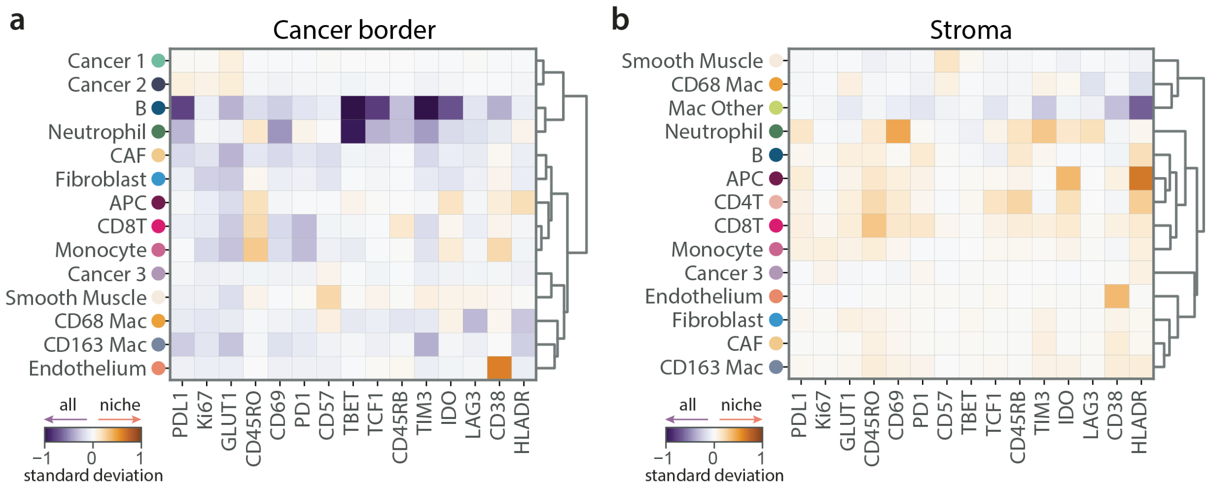


**Supplementary Figure 15: Tumor-immune border analysis.** (a) Heatmap shows the change in cell type expression within tumor border-associated niche neighborhoods as compared to all cells within the tumor border. (b) Heatmap shows the change in cell type expression within stroma-associated niche neighborhoods as compared to all cells within the stroma.

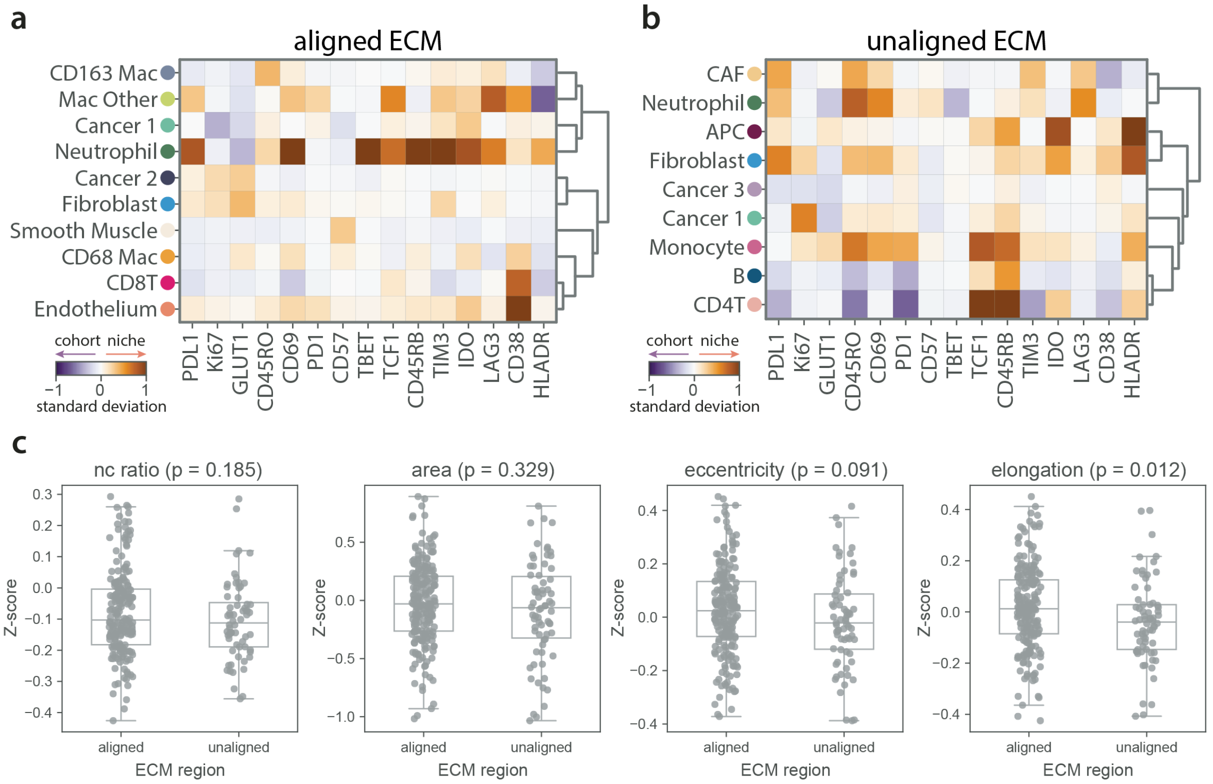


**Supplementary Figure 16: ECM alignment analysis.** (a) Heatmap shows the change in cell type expression of cellular niches within aligned extracellular matrix (ECM) regions as compared to the entire cohort. (b) Heatmap shows the change in cell type expression of cellular niches within unaligned ECM regions as compared to the entire cohort. (c) Boxplots show normalized morphological measurements of cancer cells within aligned or unaligned ECM regions. A two-sided Wilcoxon rank sum test was used to compare differences between groups.

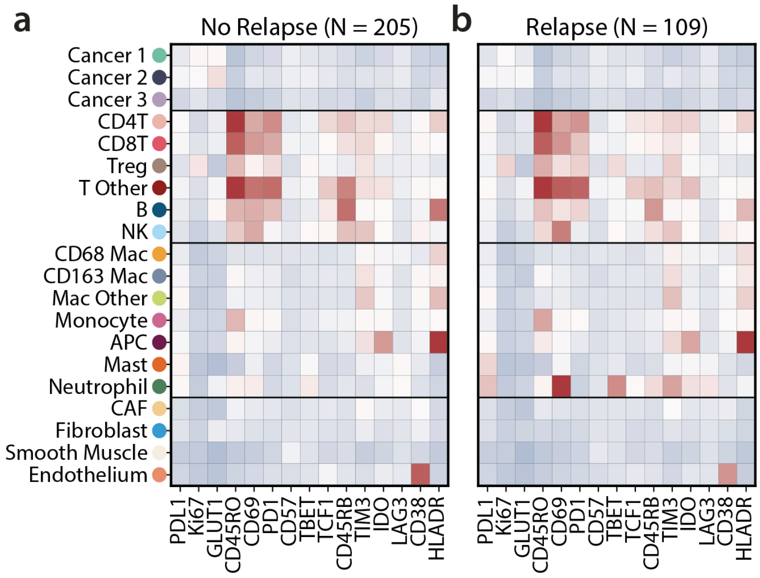


**Supplementary Figure 17: Functional expression of cellular phenotypes across recurrence status.** (a) Heatmap shows the average standardized functional expression of twenty cell phenotypes across all patients that did not relapse from the Spain TNBC cohort. (b) Heatmap shows the average standardized functional expression of twenty cell phenotypes across all patients that relapsed from the Spain TNBC cohort.

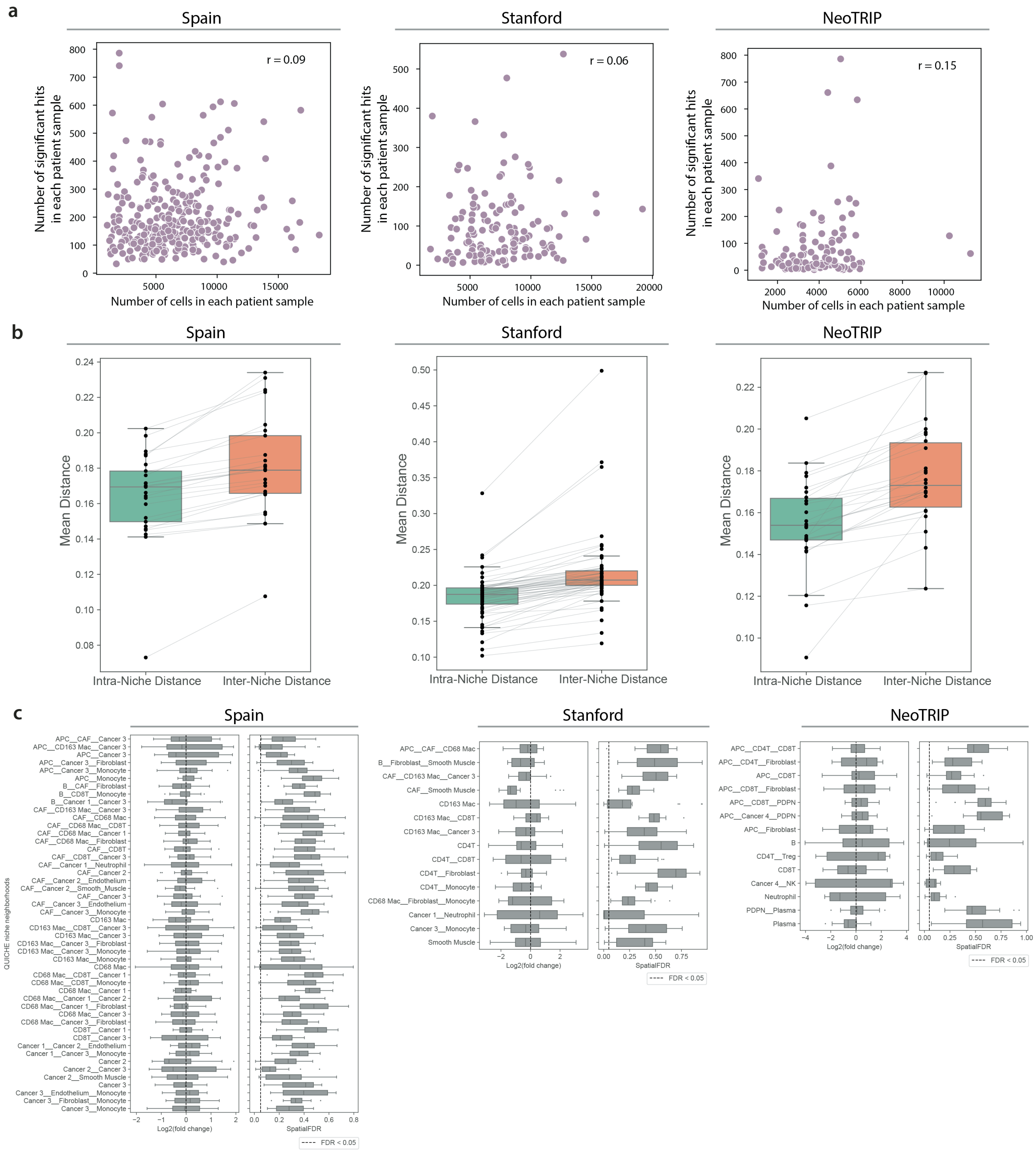


**Supplementary Figure 18: Validation of outcome-associated QUICHE analysis on TNBC cohorts.** (a) Scatter plots show the number of significant QUICHE niche neighborhoods as a function of the total number of cells in each patient sample for Spain (left), Stanford (middle), and NeoTRIP (right) TNBC cohorts. $r$ indicates the Pearson correlation coefficient. (b) Boxplots show the average distance between cells within an annotated QUICHE niche neighborhood (intra distance) as compared the remaining annotated niche neighborhoods (inter distance) in the Spain (left), Stanford (middle), and NeoTRIP (right) TNBC cohorts. Intra-distances are lower than inter-distances validating pseudobulk niche labeling performance. (c) Boxplots plots show the log2 fold change in the abundance of niche neighborhoods across permuted outcome labels (left) and spatial FDR values (right) in the Spain (left), Stanford (middle), and NeoTRIP (right) TNBC cohorts. Distributions are across 20 random trials. Log2 fold changes are centered around 0 and spatial FDR > 0.05 as expected.
